# The impact of super-spreaders in COVID-19: mapping genome variation worldwide

**DOI:** 10.1101/2020.05.19.097410

**Authors:** Alberto Gómez-Carballa, Xabier Bello, Jacobo Pardo-Seco, Federico Martinón-Torres, Antonio Salas

## Abstract

The human pathogen severe acute respiratory syndrome coronavirus 2 (SARS-CoV-2) is responsible for the major pandemic of the 21^st^ century. We analyzed >4,700 SARS-CoV-2 genomes and associated meta-data retrieved from public repositories. SARS-CoV-2 sequences have a high sequence identity (>99.9%), which drops to >96% when compared to bat coronavirus. We built a mutation-annotated reference SARS-CoV-2 phylogeny with two main macro-haplogroups, A and B, both of Asian origin, and >160 sub-branches representing virus strains of variable geographical origins worldwide, revealing a uniform mutation occurrence along branches that could complicate the design of future vaccines. The root of SARS-CoV-2 genomes locates at the Chinese haplogroup B1, with a TMRCA dating to 12 November 2019 - thus matching epidemiological records. Sub-haplogroup A2a originates in China and represents the major non-Asian outbreak. Multiple founder effect episodes, most likely associated with super-spreader hosts, explain COVID-19 pandemic to a large extent.

## Introduction

The human severe acute respiratory syndrome coronavirus 2 (SARS-CoV-2) was first detected in late 2019 in patients from the city of Wuhan (Hubei province, China) suffering from respiratory illnesses and leading to a disease that has been popularized as coronavirus disease or COVID-19. The disease was declared an International Public Health Emergency on 30 January 2020, and a few weeks later, on 11 March 2020, it was declared a pandemic by the World Health Organization ((WHO) 2020). Even though it has not been possible to trace an index case, there is a large amount of epidemiological information that has been very useful for tracking the pandemic spread of SARS-CoV-2. The first report of COVID-19 took place on 1 December 2019, in a patient from Wuhan, most likely linked to the Huanan Seafood Wholesale market of the city. Some public reports indicate that the first cases could date back to mid-November 2019. The number of cases grew gradually during December, and most of them were related to the aforementioned seafood market. In mid-January 2020, a number of patients appeared in other provinces of China, probably favored by the large annual human migration associated to the Spring Festival travel season (starting in mid-December and lasting for about 40 days). Soon covid-19 spread to other Asian countries (South Korea, 20 February 2020), and beyond: Middle East (Iran; 19 February 2020), USA (20 February 2020), Europe (Italy and Spain, 31 January 2020), etc.

Wu et al. (2020) reported the first genome sequence of SARS-CoV-2 (29,903 bp length), from a worker at the Wuhan market admitted to the Central Hospital of Wuhan on 26 December 2019; this patient experienced severe respiratory syndrome. The authors identified a new RNA virus strain belonging to the family Coronaviridae that was subsequently designated as ‘WH-Hum 1 coronavirus’ (and also ‘2019-nCoV’). According to Coutard et al. (2020) the nearest bat precursor would be RaTG13 with a genome identity to SARS-CoV-2 of 98%. Phylogenetic studies supported the theory of a natural origin for SARS-CoV-2 (Andersen et al. 2020).

Since the beginning of the COVID-19 pandemic, there has been a growing interest in exploring genetic variation in the genome of SARS-CoV-2. Identifying patterns of genomic variation can help understand the origin and spread of the pandemic and facilitate the development of future vaccines. The amount of genome data deposited in public repositories in a such a reduced timeframe offers a unique opportunity for a detailed phylogenetical characterization of SARS-CoV-2, as well as the geographic mapping of the different clades spreading worldwide, and of the impact of the outbreaks on the genome variability of the virus. Initial analyses so far used a limited number of SARS-CoV-2 genomes, and focused mostly on various evolutionary aspects of the coronavirus genomes (Andersen et al. 2020; Forster et al. 2020; Li et al. 2020; Shen et al. 2020). The Global Initiative on Sharing Avian Influenza Data GISAID (Shu and McCauley 2017) provides public access to the most complete collection of genetic sequences of several viruses, with special emphasis on influenza viruses. In 2020, the Global Initiative on Sharing Avian Influenza Data GISAID (Shu and McCauley 2017) started to compile sequence data from the virus causing COVID-19 disease, and now it makes thousands of genomic sequences of the virus available. The China National Center for Bioinformation keeps an updated resource on COVID-19 (https://bigd.big.ac.cn/ncov/tool/annotation; (Zhao et al. 2020) and provides different analytical tools to study SARS-CoV-2 variation. The open-source project Nextstrain (https://nextstrain.org; (Hadfield et al. 2018) provides an interactive web portal that allows navigating SARS-CoV-2 genome variation and helps track the spread of disease outbreaks.

In the present study, we built a solid phylogenetic skeleton of SARS-CoV-2 genomes that allows to investigate sequence variation in a large number of genomes (>4,700; **Supplementary Material**) deposited in GISAID, explore site-specific mutational instability, investigate phylogeographic patterns of variation worldwide, and clarify the role of super-spreader hosts in the pandemic.

## Results

### Identity of SARS-CoV-2 to other closely related species

Human SARS-CoV-2 genomes have a within sequence identity of 99.98% (**Table 1**); and are much more identical to bat coronavirus than to pangolin coronavirus, although the values vary substantially depending on the specimen, 93.44%–96.17% (**Table 1**). When compared to pangolin coronavirus, the range of genome identities drops to 85.24%–92.35%.

**Table 1.** Inter-specific comparisons of sequence identities between different species, including pangolin (*Manis javanica*) and bat (*Rhinolophus affinis*) against the HQ SARS-CoV-2 dataset. Comparisons involved 3,478 SARS-CoV-2 genomes against the coronaviruses indicated in the table. ID refers to identity number in GISAID (GS; omitting the prefix “EPI-ISL-“) and GenBank (GB). NC_004718.3 corresponds to the reference SARS Coronavirus genome (Marra et al. 2003). The genome #402131 corresponds to RaTG13 that has been used in the literature as bat coronavirus reference. GISAID 414518 and 420923 correspond to coronavirus analyzed from a dog and a tiger (*Panthera tigris jacksoni*) that where infected by human SARS-CoV-2. Abbreviations are as follows: Time: refers to the collection year of the specimen. Dif: Mutational differences of the coronavirus indicated when compared to the SARS-CoV-2 references sequence (MN908947.3). Id: average identity of the HQ SARS-CoV-2 against the corresponding coronavirus in the table. SD: standard deviation of Dif values. Max and Min: maximum and minimum identities shown by a SARS-CoV-2 genome with the other coronaviruses. “N” means ambiguity.

| Species | ID | Place | year | Dif | Id (%) | SD | Max | Min | C8782T | C18060T | T28144C |
| --- | --- | --- | --- | --- | --- | --- | --- | --- | --- | --- | --- |
| SARS | GB: NC_004718.3 | Toronto; Canada | 2004 | 4576 | 79.26 | 0.05 | 79.67 | 78.64 | False | True | False |
| Pangolin | GS: 410544 | Guangdong; China | 2019 | 1699 | 92.35 | 0.03 | 92.51 | 91.47 | True | False | True |
| Pangolin | GS: 410721 | Guangdong; China | 2020 | 2599 | 90.21 | 0.03 | 90.44 | 89.56 | True | False | True |
| Pangolin | GS: 412860 | China | 2019 | 2320 | 90.12 | 0.03 | 90.39 | 89.63 | True | False | True |
| Pangolin | GS: 410539 | Guangxi; China | 2017 | 3720 | 85.35 | 0.04 | 85.59 | 84.74 | True | True | True |
| Pangolin | GS: 410538 | Guangxi; China | 2017 | 3720 | 85.36 | 0.03 | 85.59 | 84.46 | True | True | True |
| Pangolin | GS: 410543 | Guangxi; China | 2017 | 3495 | 85.24 | 0.04 | 85.47 | 84.55 | True | True | N |
| Pangolin | GS: 410542 | Guangxi; China | 2017 | 3727 | 85.34 | 0.04 | 85.58 | 84.71 | True | True | True |
| Pangolin | GS: 410541 | Guangxi; China | 2017 | 3721 | 85.35 | 0.04 | 85.58 | 84.72 | True | True | True |
| Pangolin | GS: 410540 | Guangxi; China | 2017 | 3716 | 85.36 | 0.04 | 85.60 | 84.74 | True | True | True |
| Bat | GS: 402131 | Yunnan; China | 2013 | 1105 | 96.17 | 0.02 | 96.37 | 95.53 | True | True | True |
| Bat | GS: 412977 | Yunnan; China | 2019 | 1369 | 93.44 | 0.04 | 93.75 | 92.80 | True | True | True |
| Bat | GS: 412976 | Yunnan; China | 2019 | 3827 | 76.87 | 0.05 | 77.31 | 76.69 | False | False | False |
| Canine | GS: 414518 | Hong Kong | 2020 | 11 | 99.95 | 0.07 | 99.99 | 96.15 | False | False | False |
| Tiger | GS: 420293 | New York; USC | 2020 | 7 | 99.97 | 0.07 | 100.00% | 96.17 | False | False | False |
| Human | GB: MN908947.3 | Shangai; China | 2020 | 0 | 99.98 | 0.07 | 100 | 96.18 | False | False | False |

Between 1,699 and 3,727 substitution variants separate the pangolin coronavirus genomes from the SARS-CoV-2 reference sequence, and this range drops to 1,105 to 1,369 (**Table 1**) when compared to bat coronavirus. The bat #412976 coronavirus genome is conflictive because it has an unusual amount of mutational differences with respect to the SARS-CoV-2 reference and has an abnormally low sequence identity with human coronavirus (76.87%), comparable to pangolin coronavirus. This genome is problematic in the sequence alignment and should be avoided in future comparative analyses.

### Inter- and intra-specific phylogeny and the root of SARS-CoV-2

An inter-specific Maximum Likelihood (ML) tree was built using pangolin, SARS, and bat coronavirus genomes as outgroups to investigate their phylogenetic relationships with SARS-CoV-2 (**Supplementary material**). The tree depicts the SARS coronavirus genome occupying the most external branch. Next, all the pangolin genomes cluster separately from bat and human coronavirus, which also group separately. In line with its very low identity with SARS-CoV-2, bat-412976 behaves as an outlier in the tree. Overall, the clustering pattern in the tree is in very good agreement with sequence identity values (**Table 1**).

We next focused our attention on the root for all existing SARS-CoV-2 genomes, assuming the bat coronavirus as its closest coronavirus relative. We built a new ML tree including all SARS-CoV-2 genomes sequenced up to 29 February 2020 (*n* = 621); almost all of them are of Asian origin and this group should contain the Most Recent Common Ancestor (MRCA), as it is evident from phylogenetics and epidemiology that the origin of the pandemic is in China and more particularly within haplogroup B (see below and **Supplementary Material**). The ML tree unequivocally reveals that the root of SARS-CoV-2 is located in the basal B1 haplogroup (B1 genomes that do not belong to derived B1 sub-clades; **Figure 1**), and therefore points to B1 as the clade at the origin of the pandemic.

**Figure 1.**
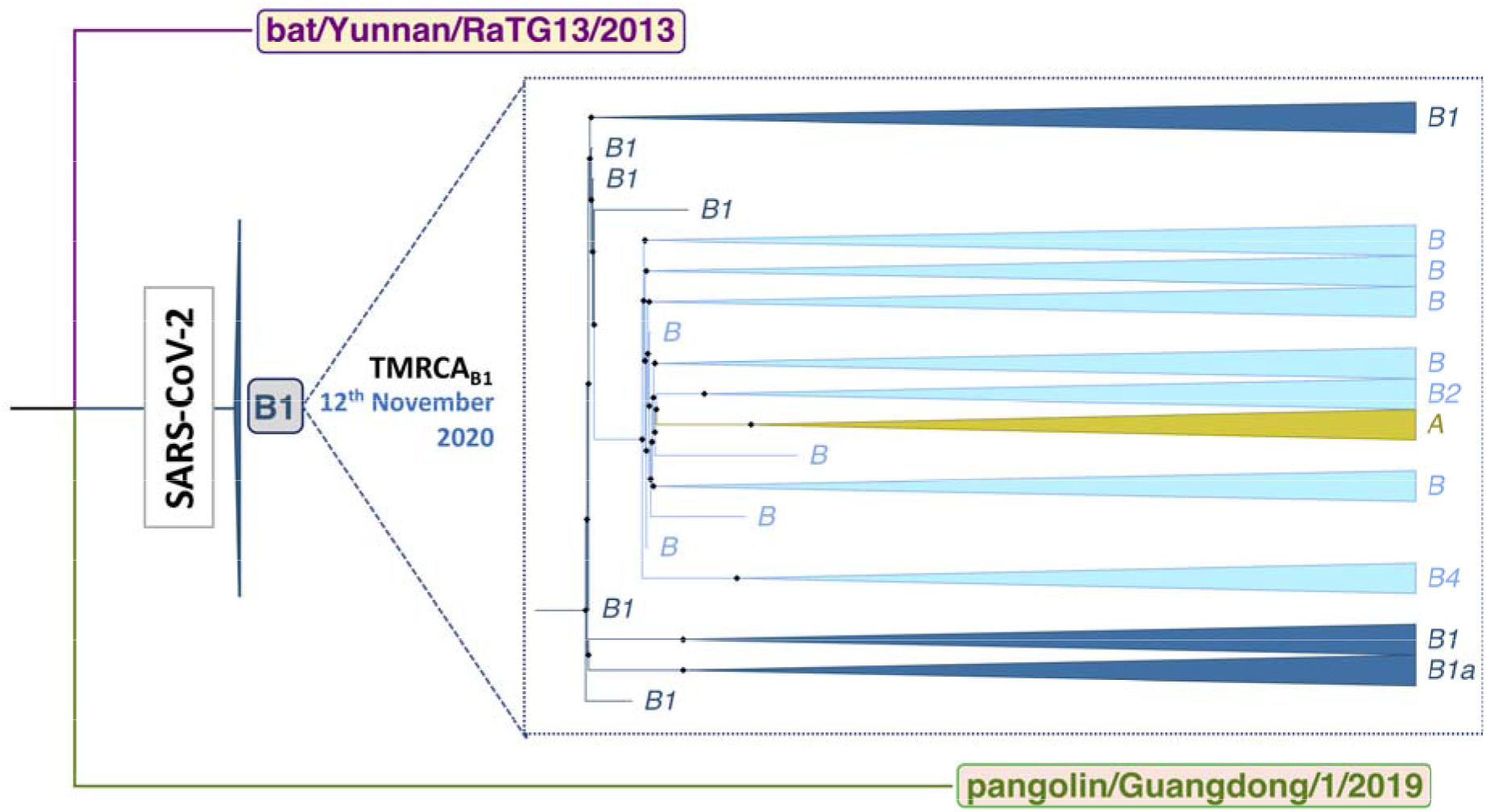
Inter-specific ML tree indicating the root of all existing SARS-CoV-2 genomes.

SARS-CoV-2 weighted mean substitution rate, as inferred from the ML tree, is 5.42×10^−4^ substitutions per site per year (s/s/y) (Bootstrap 2.5% – 75% confidence interval: 4.29×10^−4^–8.02×10^−4^ s/s/y) according to an uncorrelated relaxed-clock method; a slightly higher mutation rate of 6.05×10^−4^ s/s/y (Bootstrap 2.5% – 75% confidence interval: 4.46×10^−4^–8.22×10^−4^ s/s/y) was obtained assuming and strick-clock model.

According to a relaxed-clock model mutation rate the TMRCA for all SARS-CoV-2 genomes dates to 12^th^ November 2019 (Bootstrap 2.5% – 75% confidence interval: 7^th^ August 2019 to 8 December 2019), fully matching epidemiological dates; estimates using an strick clock mutation rate varied very little: 7^th^ November 2019 (Bootstrap 2.5% – 75% confidence interval: 18 August 2019 to 2 December 2019).

The most parsimonious tree (fully developed in **Supplementary Material Figure S4**; see also skeleton in **Figure 4A**) shows that the two very stable transitions C8782T and T28144C (3 and 1 total occurrences in the phylogeny, respectively) separate SARS-CoV-2 variation into two main clades, A and B, both originating in China. Sub-haplogroups emerging from these main clades are mainly supported by single mutations, most of them being very solid along the phylogeny (**Table S1**), and therefore granting the robustness of the different clades. It is notable that the structure of the branches in the parsimonious tree fully agrees with the skeleton shown in the ML tree.

Haplogroup B (19.65% of the genomes in the database; *n* = 664) is present in all continents; being more prevalent in North America (46.35%), South America (25.93%) and Asia (22.33%), and having the lowest frequencies in Africa (8.33%) and Europe (3.74%) (see frequency interpolated maps in **Supplementary Material**). B1 is separated from B by a single transition (C18060T) and it is by far the most numerous B subclade (*n* = 424; 63.86% of all B). The main proportion of B1 lineages worldwide is present in North America (*n* = 365; 86.04% of all existing B1 genomes). Most of the B1 genomes belong to the subclade B1a1 (B1a[A17857G]>B1a1[C1774T]); this contains at least 11 minor sub-clades, each defined by characteristic single mutations. In consonance with the root of SARS-CoV-2 being within B, we observed that basal haplogroup B is more prevalent in Asia (70%) than in anywhere else, and it is the only region containing genomes belonging to all first level B-subclades (B1, B2, B3, etc; perhaps with the exception of the minor clade B9). It is noticeable however that, within B, the fourth level sub-clade B1a1 is the most frequent haplogroup in the database (399/664; 60.09%) and it appears mainly in North America (accounting for 357 [333 in USA] out of the 399 B1a1 counts in the database; 89.47%), while it is absent in Asia; **Figure 2**. In Europe, the main B sub-clade is B3a (61.02%), which is particularly prevalent in Spain, one of the main European epicenters of COVID-19. It is most likely that most of these B3a representatives arrived in South America from Spain (where it represents 71.43% of all B genomes) given the high connectively between the two regions; **Figure 2**. The high B3 frequency observed in Spain marks a notable difference with respect to other European countries; 32 out of the 37 (86.49%) B3 genomes in Europe are located in Spain.

**Figure 2.**
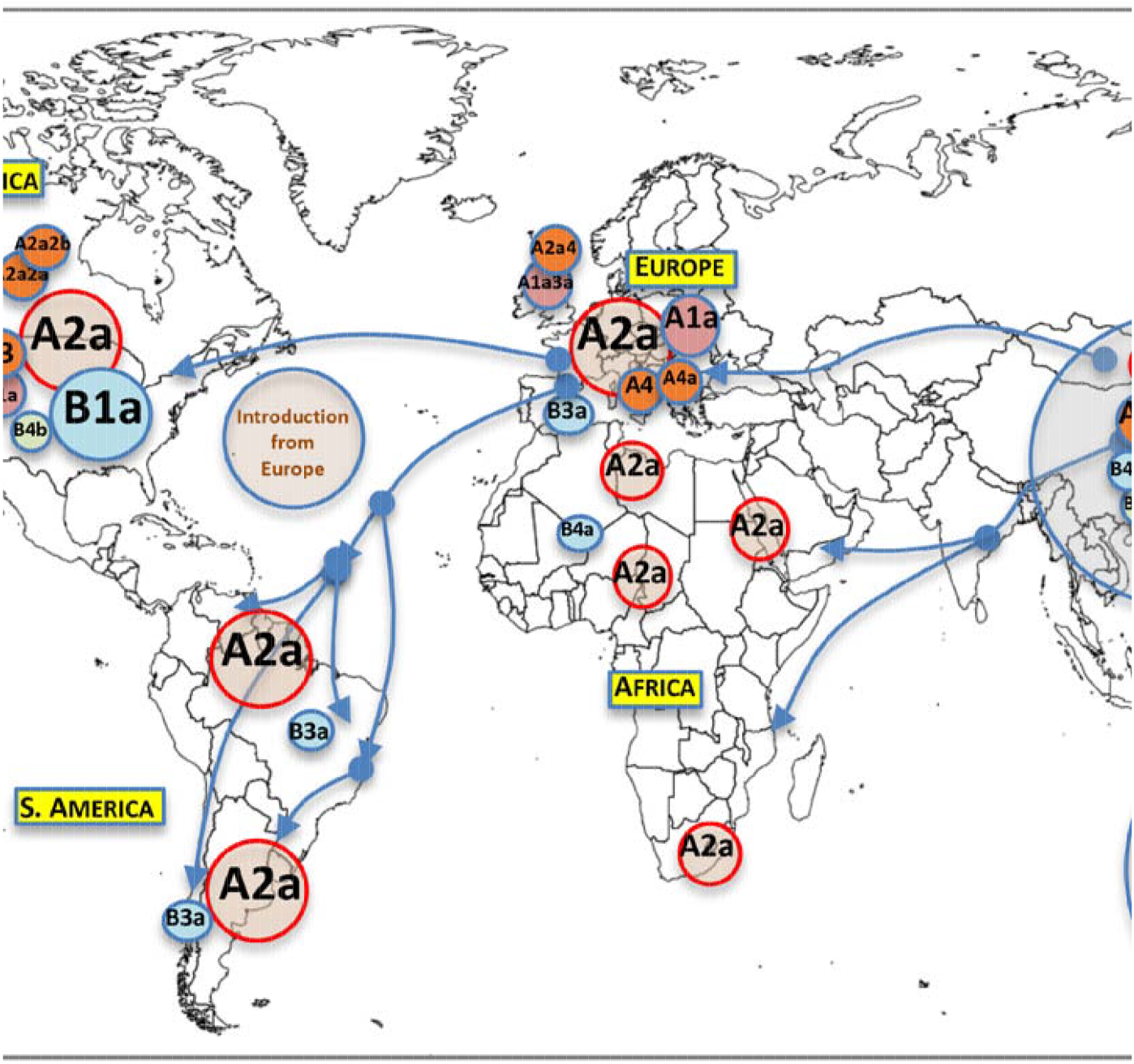
Map showing the worldwide spread of the main SARS-CoV-2 clades. Circle areas are not proportional to frequencies, and the arrows indicate just an approximate reconstruction of the phylo-dynamics of SARS-CoV-2 for the beginning of the Asian outbreak to the non-Asian spread of the pathogen based on meta-data (indicating the sampling origin and dates) and the classification of genomes into haplogroups according to the phylogeny in **Supplementary Material Figure S4**.

Haplogroup A (*n* = 2,715), with complementary frequencies to B, is the predominant clade all over the world (**Figure 2**), although with heterogeneous distributions (**Supplementary Material**). It reaches its highest frequencies in Europe (97.1%) and Africa (93.1%), is relatively high in Asia (76.3%) and Oceania (77.4%), and it has the lowest frequencies in South America (68.2%) and North America (53.1%). By far, the most frequent sub-clade of haplogroup A is A2a (*n* = 1,849; 68.10% of all A genomes), which is the main representative of the non-Asian outbreak, followed by A1a (*n* = 287; 10.57%). Even though A2a is mostly present in Europe (*n* = 1,199; 64.85% of all A2a sequences), and North America (*n* = 370; 20.01%), it most likely originated in Asia (**Supplementary Material**). A3 is mainly found in Asia, and especially in the Middle East (81.25%); its sub-clade A3a is also highly prevalent in the same region (31.94%) but shows even higher frequency in Oceania (44.44%). Other minor clades are found in more restricted areas; for instance, A4a (*n* = 39; 1.44% of A) is only found in Wales (Europe), while A5, A7 and A9b appear only in Asia.

Phylogeographic information allows reconstructing dynamics of (sub)haplogroups worldwide (**Figure 2**). The main clades emerged in Asia (mainly in China), while some minor ones appeared outside Asia (next section; **Supplementary Material**).

The number of sequences belonging to clade A and its main sub-clades increased exponentially during the outbreak occurring outside Asia at the end of February 2020, while the frequency of haplogroup B genomes increased more slowly at that time (**Supplementary Material**). Nucleotide diversity is almost homogeneous in all the different geographical regions for the main haplogroups; however, haplotype diversity (HD) values vary more substantially among haplogroups, probably indicating the weight of sequence founders in this index (see next section on super-spreaders); **Supplementary Material**.

### Super-spreaders and founder effect

It is remarkable that a few haplotypes are disproportionally represented in continental regions or in particular countries (**Supplementary Material**; **Figure S10**), appearing abruptly in a few days’ period. This pattern is compatible with super-spreaders arriving to certain geographic locations and giving rise to severe founder effects (**Figure 3A**). Haplotypes #H1, #H2, #H3, and #H4 (ID’s as in **Table S8**) are the most frequently repeated ones. H1 (*n* = 163; haplogroup A2a4), represents one of the main haplotypes responsible for the introduction or the pandemic in Europe (104/163; 63.80%), with particular frequency in the UK (35/163; 21.47%) and Belgium (23/163; 14.11%); it is also prevalent in North America (with a one week delay; 15/163 13.50%), and Australia (18/163; 11.04%). H2 (*n* = 133; A2a2a) occurs also in Europe at high frequency (75/133; 56.39%; 34 times in Iceland) and in North America (48/133; 35.82%; mostly in USA with 45 occurrences). H3 (*n* = 132; B1a1) appears at remarkably high frequency and almost exclusively in USA (126/132; 95.45%; B1a1). H4 (*n* = 78; A) corresponds to the reference sequence (GenBank acc. n° MN908947.3) and it reaches the highest frequency in Asia (60/78; 76.92%), particularly in China (49/78; 62.82%); the frequency of H4 increased in two pulses, one coinciding with its first appearance in China at the end of December 2019, and the next coinciding with the large Asian outbreak in mid-February 2020; later, H4 moved to other non-Asian locations, e.g. USA (10/78; 12.82%).

**Figure 3.**
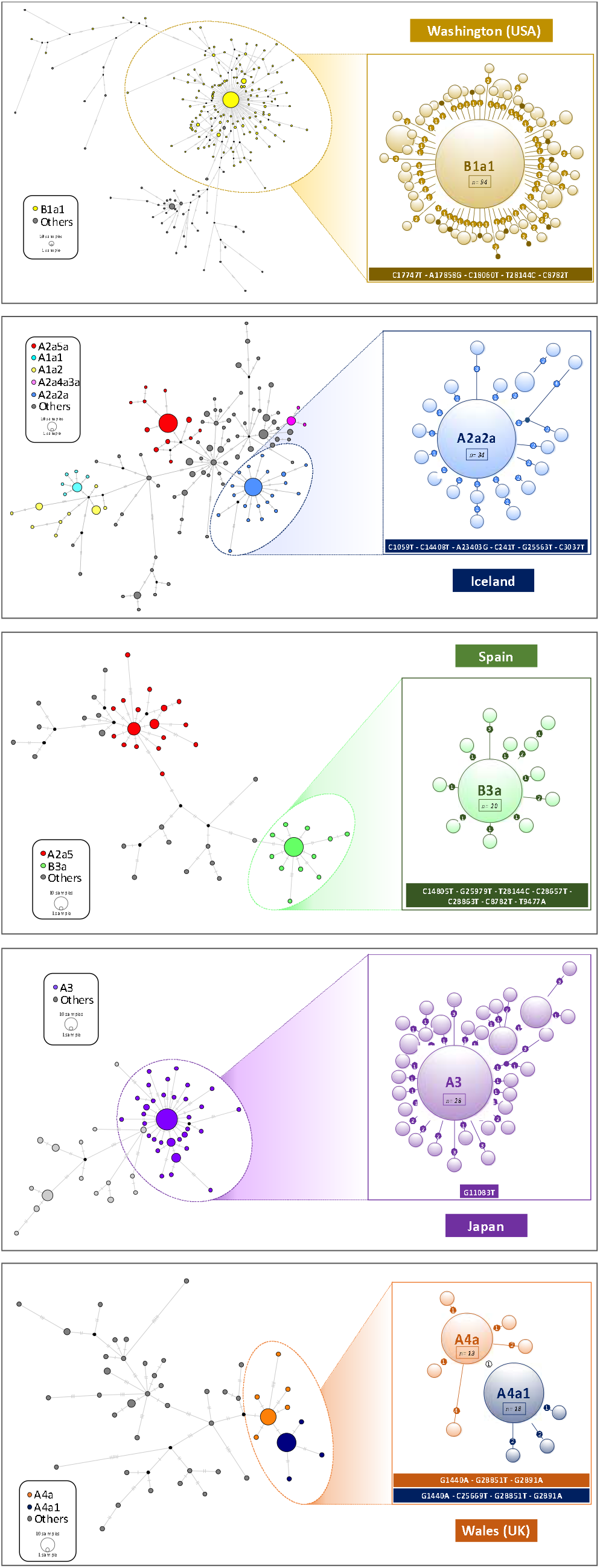
(A) Simplified SARS-CoV-2 phylogeny (see **Supplementary Material Figure S4** for the complete tree) illustrating the main branches and the main outbreaks occurring in Asia and outside Asia. The tree also shows the sub-clades that were mainly founded by a few super-spreaders. (B) EBPS based on genomes sampled from the beginning of the pandemic until the end of February 2020 (*n*=621). The orange distribution shows the real number of cases per day as recorded in https://ourworldindata.org for the same time period (we disregarded the abnormal peak occurring on 13^th^ February 2020, since more than 15,000 new cases were reported in China in just one day, most likely representing not confirmed cases); and (C) Timeline of the main events occurring during the pandemic, and indicating the MRCA of all SARS-CoV-2 genomes; the dotted area is a schematic representation of the real diversity values reported in **Supplementary Material Figure S2**. Divergence dates between SARS-CoV-2 and bat sarbecovirus reservoir and between bat and pangolin coronavirus were taken from (Boni et al. 2020).

There are additional examples of SARS-CoV-2 super-spreading (**Table S8**) appearing in restricted geographical areas. For instance, H8 (*n* = 33; A3) appears at high frequency in Japan (28/33; 84.85%). In Iceland, founder haplotypes represent a large proportion of all existing haplotypes on the island e.g. H7 exists only in Iceland (*n* = 37), and together with H2 (*n* = 34 in Iceland) and other four haplotypes, makes up 39.18% of all the haplotypes in this country. In USA, H3 occurs 126 times, and H2 45 times; together with other five haplotypes, they make up 31.75% of all genomes in this country. In the UK, eight haplotypes make-up 28.95% of the total haplotypes. H9 (*n* = 26; B3a) and H14 (*n* = 22; A2a5) are probably the main haplotypes responsible for the Spanish outbreak; H9 (21/26 in Spain; 80.77%) is particularly interesting because it belongs to haplogroup B3a, while almost all European haplotypes belong to haplogroup A (**Supplementary Material**; **Figure S9**); H14 appears 9 times in Spain (9/22; 40.91%).

Common haplotypes are frequently shared between neighboring countries, an observation mirroring the easy spread of the virus over short geographic distances; for instance, H33 (*n* = 9; of which 7 are in Portugal and 2 in Spain) or H45 (*n* = 7; of which 4 are in Portugal and 1 in Spain).

**Table 2** shows normalized phylogenetic tree features for a selected number of super-spreader candidates. For each geographical region, we additionally obtained index values for the rest of the tree excluding these candidates. The phylogenetic trees have expected values compatible with super-spreader transmissions and incompatible with super-spreader chains. For instance, clade A2a2a in Iceland has *Colless’s I* of 0.866 while it is only 0.143 when excluding candidate super-spreaders nodes. The *ILnumber* (A2a2a: 0.885 *vs*. Remaining Tree (RT): 0.415), the Sacking index (A2a2a: 0.881 *vs*. RT: 0.207), and the staircase-ness (A2a2a: 0.925 *vs*. RT: 0.685) are also consistently high as expected from super-spreader transmissions, as they are also the low values of cherries (A2a2a: 0.148 *vs*. RT: 0.592) and pitchforks (A2a2a: 0.111 *vs*. RT: 0.408). The rest of the super-spreader networks in **Table S1** follow the same pattern.

**Table 2.** Normalized phylogenetic features of super-spreader candidates. Abbreviations are as follows: *n* = total sample size; *n*_1_ = sample size of the principal node (only for super-spreader candidate); *n*_2_: sample size of derived haplotypes (only for super-spreader candidate); AL: average ladder; CH: cherries; CI: *Colless’s index;* IL: IL number; PF: pitchforks; SC: Sackin index; SN: staircase-ness.

| Clade | $n$ | $n_1$ | $n_2$ | AL | CH | CI | IL | PF | SC | SN |
| --- | --- | --- | --- | --- | --- | --- | --- | --- | --- | --- |
| Iceland – A2a2a | 54 | 34 | 20 | 0.288 | 0.148 | 0.866 | 0.885 | 0.111 | 0.881 | 0.925 |
| Iceland minus A2a5a/A1a1/A1a2/A2a4a3a/A2a2a | 117 | – | – | 0.024 | 0.592 | 0.143 | 0.415 | 0.408 | 0.207 | 0.685 |
| Japan – A3 | 69 | 28 | 41 | 0.137 | 0.174 | 0.728 | 0.851 | 0.174 | 0.757 | 0.912 |
| Japan minus A3 | 28 | – | – | 0.115 | 0.571 | 0.194 | 0.462 | 0.429 | 0.410 | 0.704 |
| Washington – B1a1 | 268 | 94 | 174 | 0.038 | 0.231 | 0.641 | 0.774 | 0.179 | 0.650 | 0.880 |
| Washington minus B1a1 | 37 | – | – | 0.067 | 0.433 | 0.389 | 0.586 | 0.250 | 0.466 | 0.763 |
| Spain – B3a | 31 | 21 | 10 | 0.897 | 0.129 | 0.933 | 0.931 | 0.097 | 0.945 | 0.933 |
| Spain minus B3a/A2a5 | 20 | – | – | 0.167 | 0.400 | 0.175 | 0.667 | 0.600 | 0.469 | 0.737 |
| Wales – A4a1 | 21 | 18 | 3 | 1.000 | 0.095 | 1.000 | 1.000 | 0.143 | 1.000 | 0.95 |
| Wales – A4a | 18 | 13 | 5 | 1.000 | 0.111 | 1.000 | 1.000 | 0.167 | 1.000 | 0.941 |
| Wales minus A4a1/A4a | 39 | – | – | 0.063 | 0.615 | 0.164 | 0.405 | 0.538 | 0.345 | 0.605 |
| Diamond Princess shipboard | 25 | 8 | 17 | 0.283 | 0.400 | 0.728 | 0.652 | 0.120 | 0.793 | 0.791 |

Networks of super-spreader candidates (**Figure 3**) show a star-like shape that is characteristic of super-spreader transmission and that clearly differentiates from patterns in the general tree that are more characteristic of homogeneous or chain of transmissions. The most outstanding super-spreader event occurred in the Washington state (**Figure 3**). This network involved about 328 genomes and its shape clearly suggests that a single super-spreader (carrying a coronavirus belonging to B1a1 lineage) could drive an important proportion of coronavirus transmissions. The data available in the present study cannot definitely identify if all these sequences in the clade were the result of one or a few super-spreader persons; such level of detail might only be determined using epidemiological and clinical data.

Values for the Diamond Princess shipboard are also comparable to those obtained for the super-spreader candidates (**Table 2**; **Supplementary Material**).Evolution of effective population size of SARS-CoV-2

Extended Bayesian Skyline Plot (EBPS) analysis undertaken on genomes sampled until the end of February (see **Supplementary Material**) reflects with great precision the main COVID-19 epidemiological episodes. If we consider the estimated TMRCA for SARS-CoV-2 to 12^th^ November 2019 and allow 14-24 days of disease incubation (until approximately the 6^th^ of December), this leaves a period of two or three weeks of silent local transmission of the virus until the first case is reported in Wuham on 30^th^ December 2019. From this moment, *N_e_* begins to slightly increase for a couple of weeks (**Figure 4B**), followed by exponential growth from 20^th^ January 2020, coinciding with the Asian outbreak. The peak is reached on 30^th^ January 2020, matching the Asian lockdown. Consequently, *N_e_* drops remarkably for the next couple of weeks, but starts to grow progressively again from 12^th^ February 2020, coinciding with the beginning of the non-Asian outbreak.

**Figure 4.**
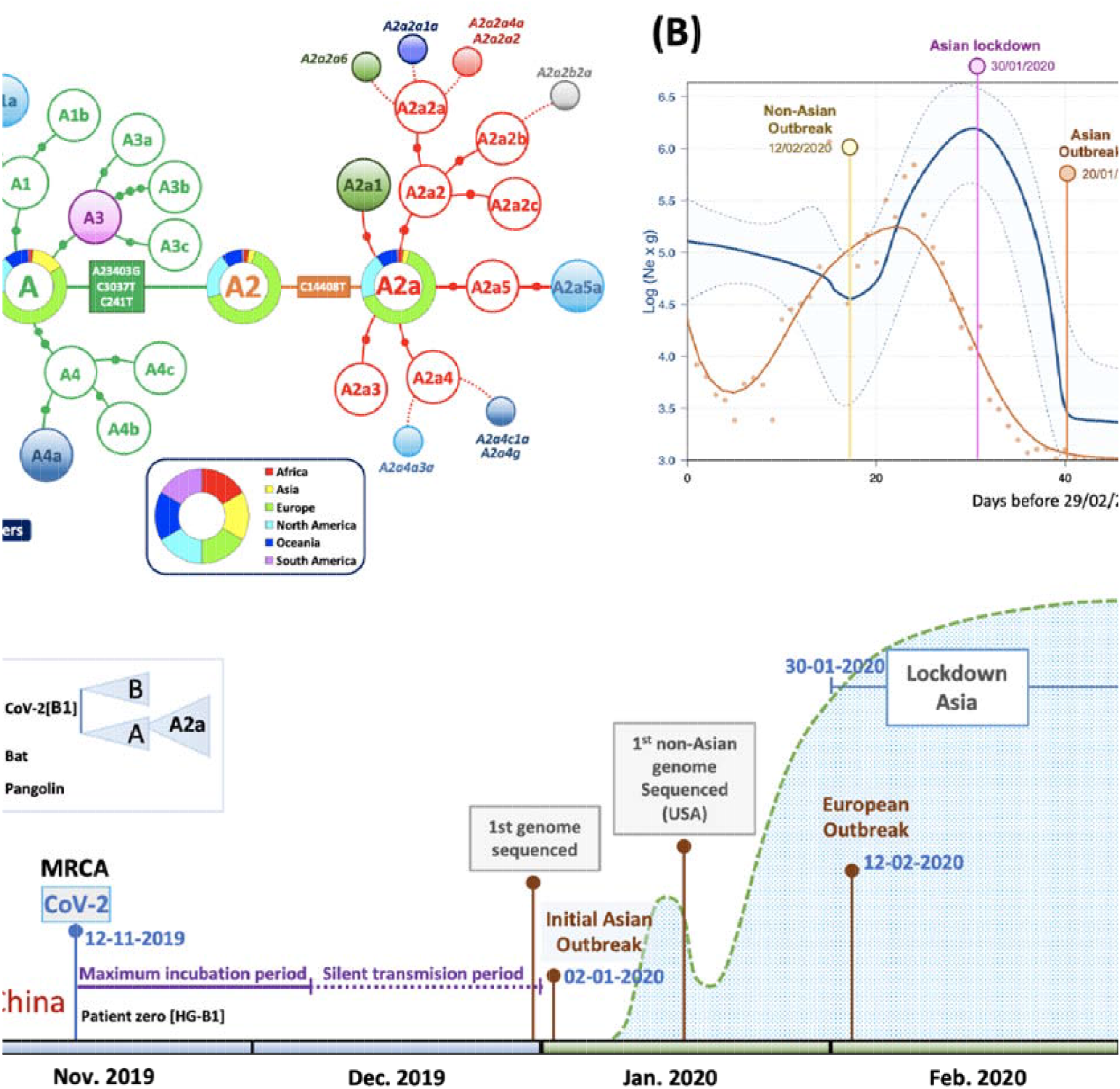

By overlapping COVID-19 incidence (officially reported cases per day worldwide; https://ourworldindata.org) with the EBPS plot, we observed comparable shape distributions, but with a remarkable 14–15 days’ delay in reported cases per day worldwide with respect to the EBPS distribution (**Figure 4B**).

## Discussion

We have undertaken a large-scale study on SARS-CoV-2 genomes considering a sample that is more than an order of magnitude higher than those of previous studies. By focusing on high-quality (HQ) genomes, we devoted great effort to elucidate the most parsimonious phylogeny of SARS-CoV-2. This effort has allowed us to present novel phylogeographic inferences on the origin and dynamics of SARS-CoV-2 strains. In particular, we discovered a few dozen genomes (representing > 1/3 of the total database) that played a fundamental role as super-spreaders of COVID-19 disease. These SARS-CoV-2 strains (belonging to different haplogroups), occur with remarkable frequency in the dataset and became founders in restricted regions or countries in short time periods (of a few days).

SARS-CoV-2 genomes show very high identity among themselves (>99%) and lower to bat coronaviruses (>96%; BatCoV RaTG13); these values are very identical to earlier estimates based on a limited number of SARS-CoV-2 genomes (Ceraolo and Giorgi 2020). The pangolin coronavirus genome, initially proposed as the original host of SARS-CoV-2, shows significantly lower identity. The high identity observed between SARS-CoV-2 genomes and other betacoronaviruses adds support to its zoonotic origin from a bat relative (Ceraolo and Giorgi 2020). The differences found between SARS-CoV-2 and their most related coronaviruses in horseshoe bat indicate that large number of mutational jumps were needed to generate these differences from a common ancestor which could have existed in a time frame between 1948-1982 (Boni et al. 2020). Divergent genomes could have been incubated in animal reservoirs before the zoonotic jump to humans in the shape of a B1 genome, in a process similar to that observed for palm civet as intermediary in other SARS coronavirus (Hu et al. 2017). These new coronaviruses would be able to use human *ACE2* receptor to infect patients. Patterns of variation observed in SARS-CoV-2 could be explained assuming a unique index case, which would already contain the very specific and well-conserved PFCS insertion. This original B1 genome would then start to diverge very soon in Wuham in two directions of the phylogeny, giving rise to its most frequent sub-lineage B1a1 and, almost simultaneously, to other B lineages and the large haplogroup A (timeline in **Figure 4C**).

According to our inferences, the TMRCA for all SARS-CoV-2 genomes would be 12^th^ November 2019. Assuming a maximum incubation time in humans of up to 24 days (Guan et al. 2020), the virus could have been infecting the first citizens from Hubei in a silent mode of transmission until the end of November 2019; and started to be noticed by Chinese health authorities in early to mid-December. The EBSP distribution suggests that the *N_e_* of SARS-CoV-2 could have started to grow significantly from 30^th^ December 2019, i.e. only two-three weeks after the initial cases reported and probably favored by super-spreaders (e.g. genomes like the reference sequence played a special role in the beginning of the Asiatic epidemic). Subsequently, it followed an exponential growth that marked the beginning of the Asian outbreak on the 20^th^ January 2020 and lasted until the end of this month. Next, *N_e_* experienced a notable drop coinciding with human intervention and quarantine implemented in Asia on 30^th^ January 2020. Finally, the beginning of a second wave of expansion outside Asia starting around 12^th^-27^th^ February 2020 is also well-recorded on the SARS-CoV-2 genomes (**Figure 4C**).

The two-week delay between the dates suggested by the EBSP distribution and the official documented incidence of COVID-19 in Asia could be due to the mean incubation time of the disease, but also to the number of cases officially declared being well below the real incidence.

With the data available in GISAID, we were not able to detect association between main haplogroups and age and sex of carriers. Further research is needed to investigate the possible differential effect of strains (haplogroups) with the disease outcome.

Evidence of natural selection acting on SARS-CoV-2 genomes needs further investigation (**Supplementary Material**), although the data suggest purifying selection acting on most of the SARS-CoV-2 genes when explored at an inter-specific level, and weaker intra-specific purifying selection. In agreement with this latter observation is the recent report indicating a 81 deletion at gene *ORF7a* that would convert the coronavirus in a less virulent pathogen with reduced short-term selective advantage (Holland et al. 2020). None of the HQ genomes investigated in our report carry this deletion.

In contrast to the weak (or null) action of positive selection on COVID-19 spread, there is strong evidence pointing to the role of genetic drift occurring in many continental regions and restricted locations, especially outside China. Phylogeographic analysis allowed us to investigate pandemic dynamics worldwide. The high incidence of a few lineages outside Asia was more probably due to drift and not selective advantages. The main non-Asian subclade A2a was probably among the first ones to leave Asia before this region established a severe population lockdown. In good agreement with epidemiological data, we observed multiple worldwide introductions of SARS-CoV-2 coming from Asia. Super-spreaders were probably the main responsible for genetic drift episodes. We detected >48 haplotypes in our dataset that most likely represent genomes transmitted by super-spreaders. Events of transmission facilitated by super-spreaders most likely occurred worldwide. In Iceland, Gudbjartsson et al. (2020) recently showed a contact tracking network (their Figure 4B) that could represent one of the many super-spreading events that existed in Iceland and that is indirectly observed in our analyses. These haplotypes have three differential features: they reached high to moderate frequencies in the population, they are characteristic of specific continental regions or even individual countries, and they appeared in a very short time period of only a few days. Network analyses of best super-spreader candidates add further support to the figure of super-spreaders in the pandemic. These analyses allowed to differentiate both haplotypes that were most likely transmitted by super-spreaders from those that were most likely transmitted by super-spreader chains, chains and homogeneous transmissions. The data suggest that these genomes have played a fundamental role in COVID-19 spreading; they alone represent 34.61% of the total genomes in the database. This finding is in good agreement with results originated from epidemiological studies and observations as well as mathematical simulations (Endo et al. 2020; Kupferschmidt 2020). The role of super-spreaders is well reported in previous pandemics, including SARS, MERS and Ebola (Stein 2011; Wong et al. 2015).

We found the 12bp polybasic furin cleavage site (PFCS) in all SARS-CoV-2 genomes with only two substitutions in two different genomes (belonging to different haplogroups; **Supplementary Material**). This segment is therefore highly mutationally stable. A BLAST search (https://blast.ncbi.nlm.nih.gov/) of the PFCS indicates that this sequence segment is absolutely specific of SARS-CoV-2. The fact that the PFCS has been found universally in all SARS-CoV-2 suggests that this insertion was acquired before the zoonotic event and not after (Andersen et al. 2020). The virulence conferred by this deletion to the coronavirus constitutes the focus of several studies (Lau et al. 2020).

The origin of SARS-CoV-2 has become a very popular question. The results of the present study (TMRCA dating of SARS-CoV-2, EBSP plot, and phylogeny) are compatible with an index case living in Wuhan-China, belonging to basal haplogroup B1, and most likely existing not before the beginning of November 2019. Subsequently, the coronavirus was transmitted from a living animal to a human host and then it started to spread from human to human. By analyzing stored biological samples from cases occurring at the beginning of the epidemy in Wuhan, it would be possible to narrow the search for patient zero among those belonging to the root of B1. The phylogeny built in the present study would be compatible with a single patient zero initiating the epidemic. Identifying the index case would help better understand how and when the spread of the pandemic begun, a lesson that would be useful in future pandemics. In agreement with previous studies (Andersen et al. 2020), the theory of SARS-CoV-2 originating artificially in a lab finds no support in the results of the present study, in the sense that variation (within and between other species), and the step-wise mutational evolution observed at SARS-CoV-2 genomes is as expected for a RNA virus in nature.

This study warrants further expansion to clarify the role of superspreaders in COVID-19 by investigating epidemiological data locally. Detecting and analyzing the genome of super-spreaders might shed light on the specific host genetic background contributing to their increased propensity to transmit the pathogen, as well as to understand the mechanisms of infection and transmission of the pathogen. Moreover, the phylogenic precision to which we classified SARS-CoV-2 genomes will also serve disease studies aimed at understanding the potential role of different pathogen strains in disease outcomes, and how these correlate to, and interact with, host genomic susceptibility.

## Material and Methods

### Database of SARS-CoV-2 sequences

We downloaded 4,721 complete genomes from the GISAID database (https://www.epicov.org/epi3/frontend) on 6 April 2020; 3,481 out of these 4,721 sequences were noted as high quality (HQ; > 29Kb, high cover only) based on the information provided by GISAID on 8 April 2020. Unless otherwise indicated, our analyses were carried out using the HQ SARS-CoV-2 genomes (see results). Although SARS-CoV-2 is an RNA virus, the data deposited in GISAID are in DNA format. Metadata for these genomes was downloaded from the Nextrain repository (https://github.com/nextstrain/ncov/tree/master/data) on 7 April 2020; this contains information on geographic location of the sample (city, country, and continental region) date of submission, age, gender, etc. This data allowed the marking of each genome by date when the virus was isolated (“Location date”) and geographic origin, among others. We also downloaded coronavirus genomes from nine pangolins and bat genomes analyzed in the Guangdong province (GISAID IDs [omitting prefix “EPI-ISL-“]): #410544, #410721, #412860, #410539, #410539, #410538, #410543, #410542, #410541, #410540), three bats (#402131, #412977, and #412976) and the reference SARS genome (GenBank acc. n°: NC_004718.3). In addition, we downloaded genomes analyzed from a tiger (GISAID: #420293) and a dog (GISAID: #414518) presumably infected with SARS-CoV-2 by humans.

The SARS-CoV-2 genomes were aligned against the reference sequence used by e.g. Nextstrain and many authors, with GenBank acc. n° MN908947.3 (submitted on 5 January 2020; GISAID ID #402125). This was the first SARS-CoV-2 genome released on GenBank.

Alignment of SARS-CoV-2 genomes against the reference sequence was carried out using MUSCLE v3.8.31 program (Edgar 2004).

Apart from the discarded low-quality (LQ) sequences, and in order to ensure comparability between genomes, we trimmed the 5’ and 3’ untranslated regions; retaining a consensus sequence of 29,607 bp that runs from position 169 to position 29,776.

### Interspecific phylogenetic analysis

We built a maximum likelihood (ML) tree in order to investigate interspecific phylogenetic relations between SARS-CoV-2 genomes and nine pangolins, three bats, and the reference SARS genome (GenBank acc. n°: NC_004718.3). Interspecific alignment was carried out using MAFFT program (Katoh et al. 2002) with default parameters. Genetic distances (F84) were computed using *dnadist* and default parameters, and the tree was built using *dnapars;* both programs are included in the software Phylip-3.697. With the SARS-CoV-2 genes aligned for all genomes with nucleotides in frame, and the ML tree, we used PAML 4 (Yang 2007) to compute the statistics ω = *Ka/Ks* (also known as *dN*/*dS*), where *Ka* is the number of non-synonymous substitutions per non-synonymous site, and *Ks* refers to synonymous substitutions per synonymous site. This ratio allows to measure the strength and mode of natural selection acting on the protein genes.

### Intraspecific phylogeny of SARS-CoV-2

There have been several attempts at reconstructing the phylogeny of SARS-CoV-2 genomes. At present there is no consensus phylogeny identifying the mutational changes characterizing main clades and sub-clades. Many of these earlier attempts used only a reduced number of genomes (Forster et al. 2020). The interactive web-phylogeny presented by Nextstrain is probably the most elaborate attempt carried out to date; this resource defines two main branches, A and B, and 9 sub-branches that cluster a variable number of sequences ranging from only 3 (haplogroup A7) or 52 sequences (haplogroup A6) to 2,279 (A2a) (28 April 2020). GISAID identifies three large clades according to changes located in the ORF8 gene and other sequence variants: a) Clade S: change L84S, with S referring to the SARS-CoV-2 spike S-glycoprotein located on the surface of the viral envelope, and sequence variant T28144C; b) Clade G: change D614G and sequence variant A23404G; and c) Clade V: NSP3-G251V and sequence variant G26144T. In addition, GISAID refers to clade ‘Other’, which is in reality a paraphyletic clade. Note that GISAID uses the reference sequence of a SARS-associated coronavirus sequenced in 2003 (Marra et al. 2003).

Here we used different strategies to build the phylogeny of SARS-CoV-2. First, a phylogeny based on ML was carried out to find the phylogenetic root of all SARS-CoV-2 using the most similar pangolin and bat coronavirus to the SARS-CoV-2 genome as ancestral relatives (GISAID IDs [omitting prefix “EPI-ISL-“]): #410721, #402131 respectively). In the particular case of SARS-CoV-2 and taking into account epidemiological evidence, we know for sure that its root is among the first genomes sequenced in China (most likely in its Hubei province). Therefore, we used only genomes available until the end of February to build the ML tree in order to reduce the noise generated by an unnecessarily large number of genomes that were sequenced later and spread outside China.

Second, the most parsimonious strategy allows to determine main and secondary sub-clades of the SARS-CoV-2 tree, identifying the characteristic mutations of clades. This phylogenetic procedure also allows to count the occurrences of mutations along branches, which serves as a good proxy for mutation-specific stability (Weissensteiner et al. 2016). We followed the HQ standards used to generate the most robust molecular phylogeny based on maximum parsimony, the human mtDNA (van Oven and Kayser 2009), although the novelty of SARS-CoV-2 genomes and the use of a variety of NGS techniques (Bandelt and Salas 2012), prevents the filtering out of sequencing errors as efficiently as in other well-known haplotypic-based phylogenies (Salas et al. 2005). Thus, we took the following steps with regards to data filtering:

1. We used only genomes labeled as ‘high-coverage only’ and ‘complete’ in GISAID.
2. We collapsed the sequences to the common sequence segment. NGS procedures generate artifacts at the 5’ and 3’ ends of the genome sequences, which generally consist of deletions. Before eliminating the extremes of the sequences, we checked in the complete genomes available if there were any variant that could be phylogenetically informative.
3. A solid phylogeny should be based on stable mutational variants; thus, only branches supported by at least five genomes were considered.
4. Since many sub-haplogroups are supported by single mutations, we only considered those having mutational stability.

We observed an excess of reversions at the tips of a few phylogenetic branches (e.g. C14805T reversion in a few A1a sequences). This phylogenetic noise could be due to the high mutational rate of the mutations involved, recombination (which is not unusual between coronaviruses (Rehman et al. 2020)), or sequencing errors. For this reason, we decided to not resolve these branches further, while we await new evidence based on higher sequence quality genomes.

By simple counting of the mutational hits along the branches and at the tips of the phylogeny, it is possible to infer the relative mutation stability of diagnostic sites. The mutations at the tips of the phylogeny were counted only once within each terminal branch. **Table S1** reports the number of occurrences in both the tree branches and in the tips of the phylogeny.

Finally, in order to guarantee the robustness of the phylogeny, we checked that inferences on the root and phylogeny estimated from the ML tree were in good agreement with the pattern of mutations observed in the most parsimonious tree.

### Phylogeny of super-spreader event

According to Colijn and Gardi (2014) *“there are simple structural properties of phylogenetic trees which, when combined, can distinguish communicable disease outbreaks with a super-spreader, homogeneous transmission and chains of transmission”*. On the other hand, Leventhal et al. (2012) states that *“The level at which a phylogenetic tree is able to resolve any contact structure depends on the rate of evolution of the pathogen. In cases such as HIV, where the rates of evolution are high enough to result in substantial genetic differences between virus populations of individual hosts, a phylogenetic tree may reveal contact structure down to the individual level”*. As the SARS-CoV-2 has a comparable mutation rate as HIV (Lemey et al. 2006; Patiño-Galindo and González-Candelas 2017; Zanini et al. 2017), we carried out phylogenetic analysis aimed at distinguishing patterns of super-spreader transmission and super-spreader chain of transmission.

In particular, we are interested in computing phylogenetic features of SARS-CoV-2 transmissions for our best candidates (those haplotypes that have experienced a high frequency locally and in a short time period) and compare them to those obtained using the remaining haplotypes from the same specific region. We first built phylogenetic trees from sequence alignments using *SplitsTree5* (Huson and Bryant 2006). We used the R library *phylotop* (Kendall et al. 2018) to calculate the following tree features (Colijn and Gardy 2014):

- *Cherry number/n:* It is the number of cherries over number of leaves. It is slightly lower and more variable for super-spreader out-breaks.
- *Colless’s I* (index) or imbalance: It reaches higher values for superspreader out-breaks
- *IL number*: portion of internal nodes with one leaf descendant. It reaches higher values for super-spreader out-breaks
- *Pitchforks:* as cherries, pitchforks decrease with increasing infection rate (Metzig et al. 2019).
- *Sacking index:* it is the mean path length form tip to root. Slightly higher in super-spreader outbreaks and lower in chain out-breaks.
- *Staircase-ness:* defined as the portion of imbalance nodes (Norström et al. 2012). It shows much lower values for chain out-breaks.

These values were computed for the main super-spreader candidates in specific geographic regions and also for the tree that includes the remaining haplotypes from the same region. The comparison between values in both trees allows to compare different transmission patterns from the same cultural/social context.

In addition, we also carried out network analyses for a reported event of a super-spreader occurred in the Diamond Princess shipboard for which we could access to 25 SARS-CoV-2 genomes. According to Sekizuka et al. (2020) the super-spreading event occurring in the shipboard may have originated from a single COVID-19 patient who had disembarked in Hong Kong on 25 January 2020.

To visualize the super-spreader phylogeny, we built phylogenetic median joining networks using *POPART* software (Leigh and Bryant 2015).

### Statistical analysis

The average number of nucleotide differences per site between DNA sequences or nucleotide diversity (π) (Nei and Li 1979), sequence/haplotype diversity (HD) and Tajima’s *D* statistics (Tajima 1989) were computed for the main continental regions, haplogroups and gene partitions. Tajima’s *D* is a test for neutrality in the context of infinite-sites model of sequence evolution and it is negligibly affected by *S*, sample size, and recombination (Ramirez-Soriano et al. 2008).

Multidimensional Scaling (MDS) was undertaken to identify clusters of genetic variation by examining (a) all the variation observed in the SARS-CoV-2 genomes and (b) the phylogenetic diagnostic variants of the SARS-CoV-2 tree inferred from parsimony. For this, we used the function *cmdscale* (library *stats*) from the statistical software R Project for Statistical Computing v. 3.3.1 (https://www.r-project.org/; (Team 2012)).

The geographic representation of haplogroup frequencies in world maps was carried out using SAGA v. 7.6.2 (http://www.saga-gis.org/) (Conrad et al. 2015), and the ordinary Kriging method. We used only sampling points with 10 samples or more; to avoid unnecessary loss of sampling points, a few of them were collapsed into nearest points to represent local areas, whenever possible, in order to reach the minimum sampling required.

From the sequence alignments and annotated files, we summarized information on mutational patterns in the SARS-CoV-2 genomes (**Table S2 and S3**). We used a Fisher’s exact test to check if there are differences in the transition-to-transversion ratio (ts/tv), and the synonymous/non-synonymous changes.

We carried out the ML phylogeny and Extended Bayesian Skyline plot analysis using the sequences collected before 29 February 2020 (*n* = 621). We constructed the ML tree using RAxML-HPC v.8 (Stamatakis 2014) using rapid bootstrapping analysis with 1,000 iterations. The best maximum likelihood tree was visualized and edited using FigTree v. 1.4.4 (http://tree.bio.ed.ac.uk/software/figtree/). The ML tree and sampling dates were used for estimating the Time of the Most Recent Common Ancestor (TMRCA) and molecular rates, fitting a molecular clock to the phylogeny through a fast relaxed-clock method based on a Gamma-Poisson mixture model of substitution rates, and using the R package *treedater* (Volz and Frost 2017). We selected the best molecular clock model by testing if the relaxed-clock offers better fit to data, and also identified and removed tip outlier lineages that fit poorly the substitution model in order to obtain a tree that could fit better the molecular clock. We estimated confidence intervals for rates and dates using a parametric bootstrap approach.

The demography of SARS-CoV-2 sequences was inferred using the Extended Bayesian Skyline Plot method (EBSP) (Heled and Drummond 2008) implemented in BEAST v2.6.2 (Drummond and Rambaut 2007). EBSPs allow the inference of effective population size (N_*e*_) through time and also estimate the number of demographic changes from the data. We used strict-clock and a rate of evolution of 0.80 x 10^−3^ [0.14 x 10^−3^ – 1.31 x 10^−3^] s/s/y based on recent estimations (http://virological.org/t/phylodynamic-analysis-90-genomes-12-feb-2020/356). Two independent Markov chain-Monte Carlo runs of 200,000,000 steps each were performed, with samples taken every 1,000 steps and 10% discarded as burn-in. Following Tracer (v. 1.6) output (http://tree.bio.ed.ac.uk/software/tracer/) (Drummond and Rambaut 2007) inspection for distributions convergence, both runs were combined independently using LogCombiner v1.8.2 (Drummond and Rambaut 2007), with 10% discarded as burn-in. EBPS data was plotted using R software (Team 2012).

The meta-data in GISAID contains information on gender and age of the COVID-19 patients. This data, in combination with the phylogeny, allows investigating for possible association among haplogroups, gender and age. Association tests were carried out for the main (sub)haplogroups (A, B, A1a, A2a, A3, A3a, A4a, A9b, B1a, B3a; all with sample size > 50). We carried out a Mann-Whitney test to analyze haplogroup association with age. Because each region has its own haplogroup frequency patterns and epidemiological characteristics, we considered a non-parametric Kruskal-Wallis test using all the sampling data per regions and main haplogroups. Association between haplogroups/regions and gender was explored using the Fisher’s exact test. The nominal significant value was set to 0.05. Bonferroni adjustment was used to account for multiple testing.

In-house R and Perl (http://www.perl.org) scripts were used to display results obtained from the different software packages used.

## Acknowledgements

This study received support from the Instituto de Salud Carlos III: project GePEM (Instituto de Salud Carlos III(ISCIII)/PI16/01478/Cofinanciado FEDER), DIAVIR (Instituto de Salud Carlos III(ISCIII)/DTS19/00049/Cofinanciado FEDER; Proyecto de Desarrollo Tecnológico en Salud) and Resvi-Omics (Instituto de Salud Carlos III(ISCIII)/PI19/01039/Cofinanciado FEDER) and project BI-BACVIR (PRIS-3; Agencia de Conocimiento en Salud (ACIS)—Servicio Gallego de Salud (SERGAS)—Xunta de Galicia; Spain) given to A.S.; and project ReSVinext (Instituto de Salud Carlos III(ISCIII)/PI16/01569/Cofinanciado FEDER), and Enterogen (Instituto de Salud Carlos III(ISCIII)/PI19/01090/Cofinanciado FEDER) given to F.M.-T.

We gratefully acknowledge GISAID and contributing laboratories for giving us access to the SARS-CoV-2 genomes used in the present study.

## Notes

### Competing Interest Statement

The authors have declared no competing interest.

### Summary of Updates

v.2. Material and Methods section has been moved into the body text. We have updated ORCID to the rest of the authors. We have corrected the name of one of the authors into the bioRxiv system (Xabi should be Xabier). v.3. We have now added phylogenetic network analysis on best super-spreader candidates and statistical indices favoring the figure of super-spreaders in the COVID-19 pandemic.

## References

(WHO) WHO. 2020. WHO Director-General’s opening remarks at the media briefing on COVID-19 – 11 March 2020.

Andersen KG, Rambaut A, Lipkin WI, Holmes EC, Garry RF. 2020. The proximal origin of SARS-CoV-2. Nat Med 26: 450–452.

Bandelt HJ, Salas A. 2012. Current next generation sequencing technology may not meet forensic standards. Forensic Sci Int Genet 6: 143–145.

Boni MF, Lemey P, X J, T.T.-Y. L, Perry B, Castoe T, Rambaut A, Robertson DL. 2020. Evolutionary origins of the SARS CoV 2 sarbecovirus lineage responsible for the COVID-19 pandemic. bioRxiv doi: https://doi.org/10.1101/2020.03.30.015008.

Ceraolo C, Giorgi FM. 2020. Genomic variance of the 2019-nCoV coronavirus. Journal of medical virology 92: 522–528.

Colijn C, Gardy J. 2014. Phylogenetic tree shapes resolve disease transmission patterns. Evol Med Public Health 2014: 96–108.

Conrad O, Bechtel B, Bock M, Dietrich H, Fischer E, Gerlitz L, Wehberg J, Wichmann V, Böhner J. 2015. System for automated geoscientific analyses (SAGA) v. 2.1. 4. Geoscientific Model Development 8: 1991–2007.

Coutard B, Valle C, de Lamballerie X, Canard B, Seidah NG, Decroly E. 2020. The spike glycoprotein of the new coronavirus 2019-nCoV contains a furin-like cleavage site absent in CoV of the same clade. Antiviral Res 176: 104742.

Drummond AJ, Rambaut A. 2007. BEAST: Bayesian evolutionary analysis by sampling trees. BMC Evol Biol 7: 214.

Edgar RC. 2004. MUSCLE: multiple sequence alignment with high accuracy and high throughput. Nucleic Acids Res 32: 1792–1797.

Endo A, Abbott S, Kucharski AJ, Funk S. 2020. Estimating the overdispersion in COVID-19 transmission using outbreak sizes outside China. Wellcome Open Res 5: 67.

Forster P, Forster L, Renfrew C, Forster M. 2020. Phylogenetic network analysis of SARS-CoV-2 genomes. Proc Natl Acad Sci U S A doi:10.1073/pnas.2004999117.

Guan WJ, Ni ZY, Hu Y, Liang WH, Ou CQ, He JX, Liu L, Shan H, Lei CL, Hui DSC et al. 2020. Clinical Characteristics of Coronavirus Disease 2019 in China. N Engl J Med doi:10.1056/NEJMoa2002032.

Gudbjartsson DF, Helgason A, Jonsson H, Magnusson OT, Melsted P, Norddahl GL, Saemundsdottir J, Sigurdsson A, Sulem P, Agustsdottir AB et al. 2020. Spread of SARS-CoV-2 in the Icelandic Population. N Engl J Med doi: 10.1056/NEJMoa2006100.

Hadfield J, Megill C, Bell SM, Huddleston J, Potter B, Callender C, Sagulenko P, Bedford T, Neher RA. 2018. Nextstrain: real-time tracking of pathogen evolution. Bioinformatics 34: 4121–4123.

Heled J, Drummond AJ. 2008. Bayesian inference of population size history from multiple loci. BMC Evol Biol 8: 289.

Holland LA, Kaelin EA, Maqsood R, Estifanos B, Wu LI, Varsani A, Halden RU, Hogue BG, Scotch M, Lim ES. 2020. An 81 nucleotide deletion in SARS-CoV-2 ORF7a identified from sentinel surveillance in Arizona (Jan-Mar 2020). Journal of virology doi: 10.1128/JVI.00711-20.

Hu B, Zeng LP, Yang XL, Ge XY, Zhang W, Li B, Xie JZ, Shen XR, Zhang YZ, Wang N et al. 2017. Discovery of a rich gene pool of bat SARS-related coronaviruses provides new insights into the origin of SARS coronavirus. PLoS pathogens 13: e1006698.

Huson DH, Bryant D. 2006. Application of phylogenetic networks in evolutionary studies. Mol Biol Evol 23: 254–267.

Katoh K, Misawa K, Kuma K, Miyata T. 2002. MAFFT: a novel method for rapid multiple sequence alignment based on fast Fourier transform. Nucleic Acids Res 30: 3059–3066.

Kendall M, Boyd M, Colijn C. 2018. Calculating topological properties of phylogenies.

Kupferschmidt K. 2020. Why do some COVID-19 patients infect many others, whereas most don’t spread the virus at all?

Lau SY, Wang P, Mok BW, Zhang AJ, Chu H, Lee AC, Deng S, Chen P, Chan KH, Song W et al. 2020. Attenuated SARS-CoV-2 variants with deletions at the S1/S2 junction. Emerging microbes & infections 9: 837–842.

Leigh JW, Bryant D. 2015. POPART : full-feature software for haplotype network construction. Methods Ecol Evol: 1110–1116.

Lemey P, Rambaut A, Pybus OG. 2006. HIV evolutionary dynamics within and among hosts. AIDS Rev 8: 125–140.

Leventhal GE, Kouyos R, Stadler T, Wyl V, Yerly S, Boni J, Cellerai C, Klimkait T, Gunthard HF, Bonhoeffer S. 2012. Inferring epidemic contact structure from phylogenetic trees. PLoS computational biology 8: e1002413.

Li X, Zai J, Zhao Q, Nie Q, Li Y, Foley BT, Chaillon A. 2020. Evolutionary history, potential intermediate animal host, and cross-species analyses of SARS-CoV-2. J Med Virol doi:10.1002/jmv.25731.

Marra MA, Jones SJ, Astell CR, Holt RA, Brooks-Wilson A, Butterfield YS, Khattra J, Asano JK, Barber SA, Chan SY et al. 2003. The Genome sequence of the SARS-associated coronavirus. Science 300: 1399–1404.

Metzig C, Ratmann O, Bezemer D, Colijn C. 2019. Phylogenies from dynamic networks. PLoS Comput Biol 15: e1006761.

Nei M, Li WH. 1979. Mathematical model for studying genetic variation in terms of restriction endonucleases. Proc Natl Acad Sci U S A 76: 5269–5273.

Norström MM, Prosperi MC, Gray RR, Karlsson AC, Salemi M. 2012. PhyloTempo: A Set of R Scripts for Assessing and Visualizing Temporal Clustering in Genealogies Inferred from Serially Sampled Viral Sequences. Evol Bioinform Online 8: 261–269.

Patiño-Galindo JA, González-Candelas F. 2017. The substitution rate of HIV-1 subtypes: a genomic approach. Virus Evol 3: vex029.

Ramirez-Soriano A, Ramos-Onsins SE, Rozas J, Calafell F, Navarro A. 2008. Statistical power analysis of neutrality tests under demographic expansions, contractions and bottlenecks with recombination. Genetics 179: 555–567.

Rehman SU, Shafique L, Ihsan A, Liu Q. 2020. Evolutionary Trajectory for the Emergence of Novel Coronavirus SARS-CoV-2. Pathogens 9.

Salas A, Carracedo A, Macaulay V, Richards M, Bandelt HJ. 2005. A practical guide to mitochondrial DNA error prevention in clinical, forensic, and population genetics. Biochem Biophys Res Commun 335: 891–899.

Sekizuka T, Itokawa K, Kageyama T, Saito S, Takayama I, Asanuma H, Nao N, Tanaka R, Hashino M, Takahashi T et al. 2020. Haplotype networks of SARS-CoV-2 infections in the Diamond Princess cruise ship outbreak. medRxiv doi:https://doi.org/10.1101/2020.03.23.20041970.

Shen Z, Xiao Y, Kang L, Ma W, Shi L, Zhang L, Zhou Z, Yang J, Zhong J, Yang D et al. 2020. Genomic diversity of SARS-CoV-2 in Coronavirus Disease 2019 patients. Clin Infect Dis doi:10.1093/cid/ciaa203.

Shu Y, McCauley J. 2017. GISAID: Global initiative on sharing all influenza data - from vision to reality. Euro Surveill 22.

Stamatakis A. 2014. RAxML version 8: a tool for phylogenetic analysis and post-analysis of large phylogenies. Bioinformatics 30: 1312–1313.

Stein RA. 2011. Super-spreaders in infectious diseases. International journal of infectious diseases : IJID : official publication of the International Society for Infectious Diseases 15: e510–513.

Tajima F. 1989. Statistical method for testing the neutral mutation hypothesis by DNA polymorphism. Genetics 123: 585–589.

Team TRC. 2012. R: A Language and Environment for Statistical Computing. (ed. RCT The).

van Oven M, Kayser M. 2009. Updated comprehensive phylogenetic tree of global human mitochondrial DNA variation. Hum Mutat 30: E386–E394.

Volz EM, Frost SDW. 2017. Scalable relaxed clock phylogenetic dating. Virus Evol 3: vex025.

Weissensteiner H, Pacher D, Kloss-Brandstätter A, Forer L, Specht G, Bandelt H-J, Kronenberg F, Salas A, Schonherr S. 2016. HaploGrep 2: mitochondrial haplogroup classification in the era of high-throughput sequencing. Nucleic acids research 44: W58–W63.

Wong G, Liu W, Liu Y, Zhou B, Bi Y, Gao GF. 2015. MERS, SARS, and Ebola: The Role of Super-Spreaders in Infectious Disease. Cell Host Microbe 18: 398–401.

Wu F, Zhao S, Yu B, Chen YM, Wang W, Song ZG, Hu Y, Tao ZW, Tian JH, Pei YY et al. 2020. A new coronavirus associated with human respiratory disease in China. Nature 579: 265–269.

Yang Z. 2007. PAML 4: phylogenetic analysis by maximum likelihood. Mol Biol Evol 24: 1586–1591.

Zanini F, Puller V, Brodin J, Albert J, Neher RA. 2017. In vivo mutation rates and the landscape of fitness costs of HIV-1. Virus Evol 3: vex003.

Zhao WM, Song SH, Chen ML, Zou D, Ma LN, Ma YK, Li RJ, Hao LL, Li CP, Tian DM et al. 2020. The 2019 novel coronavirus resource. Yi Chuan 42: 212–221.

