## Supplementary Material for "The impact of super-spreaders in COVID-19: mapping genome variation worldwide"

**INDEX**

**Supplementary Results**

The SARS-CoV-2 genome database

Molecular variation of SARS-CoV-2 genomes

The root of SARS-CoV-2 genomes

Intraspecific phylogeny of SARS-CoV-2

Considerations on the parsimonious phylogeny

Further considerations on phylogeographic patterns of SARS-CoV-2 genomes

Clade A2 and the main outbreak outside Asia

Considerations on natural selection acting on SARS-CoV-2 genomes

Further considerations on super-spreaders and founder effect

Association test of haplogroups with sex and age

**References**

**Supplementary Results**

**The SARS-CoV-2 genome database**

We initially investigated 4,716 SARS-CoV-2 genomes; a proportion of them (71.9%; *n* = 3,393) were identified by GISAID as having “<1% of NNN” and complete (referred herein as the HQ dataset). We first explored the differences between the full dataset and the HQ sequences. An indirect indication of the quality of the sequences is the sequence length, which is on average 25,636 bp for the HQ dataset, compared to a significantly lower value for the full database, namely 23,736 bp. Moreover, the average number of ambiguities (including all kinds of IUPAC codes) per sequence in the full database is 6.5x the amount of ambiguities in the HQ database (249.3 *versus* 37.9). The large number of ambiguities and indels existing in the LQ distorts sequence alignment and, consequently, a correct sequence annotation and phylogenetic reconstruction. Still, we counted an average of 43.5 ambiguities per genome in the HQ dataset. It is also important to mention that different platforms were used to sequence SARS-CoV-2 genomes, and corresponding differences between them are evident when examining the processed GISAID files (**Figure S1**). We examined several quality variables in the genomes sequenced using different NGS technologies, by taking 60 random genomes from GISAID, respectively 30 of LQ and HQ (information on quality cannot be datamined from GISAID but has to be taken manually). The number of N’s islands and the length of NNN’s stretches are significantly higher in the LQ dataset, while the length of the genomes is higher in the HQ genomes after eliminating the ambiguities; the latter is partly due to the much higher number of ambiguities existing in the 5’ and 3’ ends in the LQ datasets. When using the LQ filter, Illumina yields a significant higher number of N’s compared to Nanopore, but these differences disappear when looking at the HQ set (**Figure S1**). Indels are more common in the HQ dataset, and Nanopore seems to capture more than Illumina.

In order to minimize the effect caused by potential sequencing errors, only the HQ genomes were used for the subsequent analysis. In addition to LQ sequences, we also eliminated fifty-three of the HQ sequences lacking sampling date or having ambiguous haplogroup adscription (see below). Thereby, the final number of HQ genomes used for all the analyses was 3,340 (1,725 unique sequences; 51.57%).

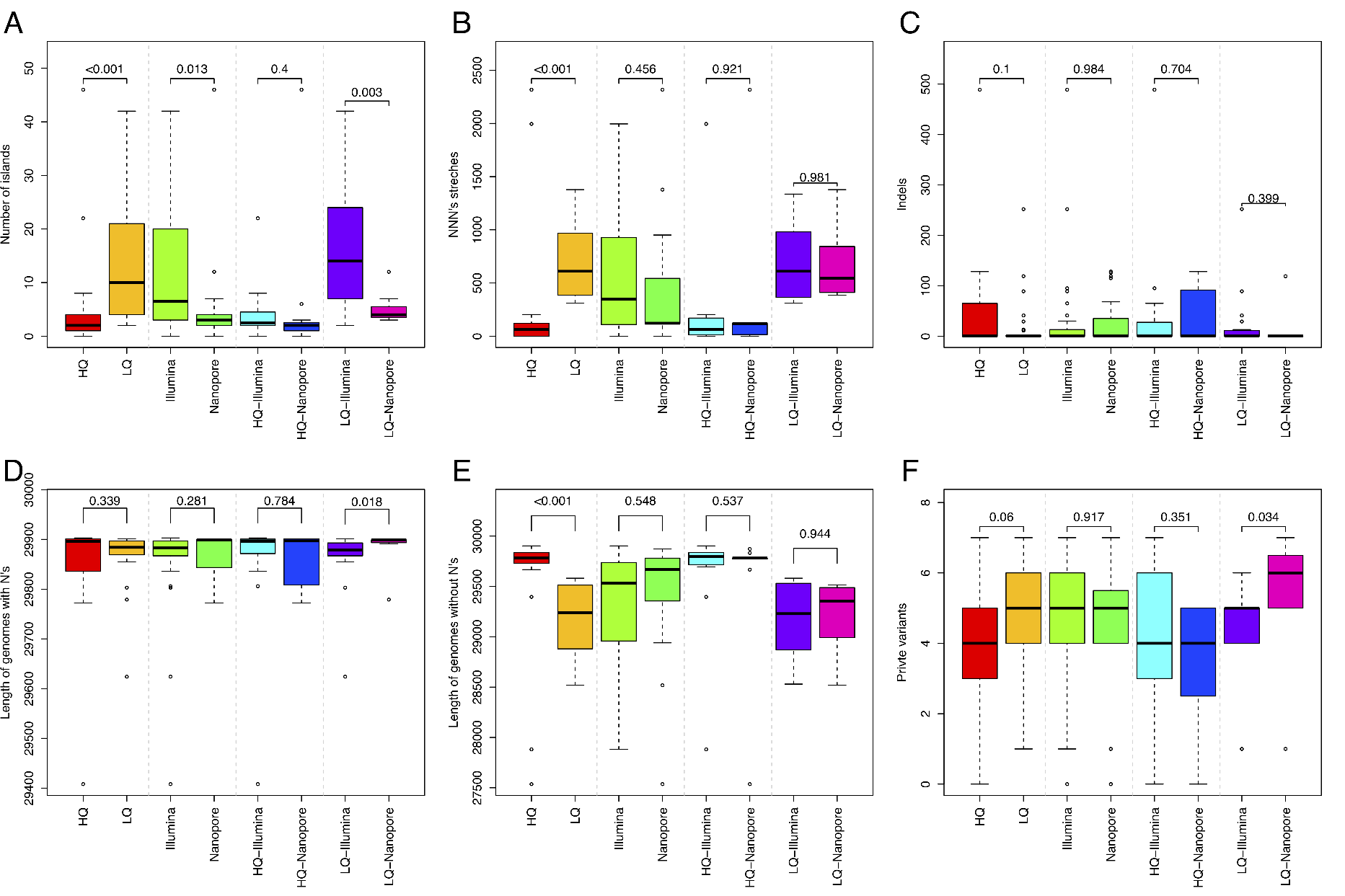

**Figure S1**. A total of 30 LQ and 30 HQ sequences were downloaded from GISAID, with information on their NGS technology used retrieved manually from the database. Comparisons between technologies were carried out on the FASTA files already processed by GISAID.

**Molecular variation of SARS-CoV-2 genomes**

We identified a large number of indels in our dataset: 2,209 insertions and 25 deletions. We also counted 2,159 substitutions and 49 Multi-nucleotide Position Polymorphisms (MNPs). From the data available, it is not possible to reconstruct the mutational process leading to MNP variants; for instance, GGG28881AAC could be interpreted as a single mutational event (MNPs) or as the concatenation of three independent mutational events: G28881A+G28881A+G28881C. Here we opted for the most parsimonious solution, namely considering them as single mutational events (in contrast to e.g. CNCB, which favors the latter option). Although the parsimonious interpretation entails the risk of under-estimating overall and allele-specific mutation rates, the few MNPs existing would only elevate the number of mutations from 49 to 195 events. In this regard, there is evidence to suggest that mutations in MNPs are not independent (*1*). Indels and MNPs were disregarded from most of the analyses.

The substitutions observed in SARS-CoV-2 genomes show the following pattern: (a) transitions (*n* = 1,524), with 72% being purine transitions (C <–> T) and 28% pyrimidine transitions (A <–> G); and (b) transversions (*n* = 644), with 76% being purine to pyrimidine, and 24% pyrimidine to purine. The ts/tv ratio is 2.37 (**Table S2**). In comparisons between humans and chimpanzees, Ebersberger et al. (*2*) found a *ts*/*tv* ratio of 2.4 – in agreement with our ts/tv ratio. Further, according to DePristo et al. (*3*), the ts/tv ratio is expected to be 2.1 for whole-genome sequencing and 2.6-3.3 for exome sequencing in human studies, while Zook et al. (*4*) have stated that very low ratios (0.5) generally point to sequence artifacts. Therefore, our ratio falls within the expected range for a good quality dataset in terms of substitutions. The ts/tv ratio was similar when considering the main clades of SARS-CoV-2 (according to the phylogeny described below); namely, 2.38 for haplogroup A, 2.74 for haplogroup B, and 2.42 for haplogroup A2a. We also investigated the pattern of mutations by genes (**Table S3**). The highest *ts*/*tv* ratio occurs in gene E (4.33) and *ORF1ab* (3.05), and the lowest in gene *ORF6* (1.11) and the intergenic regions (1.03); these differences are however not statistically significant under a Fisher’s exact test, and considering an strict Bonferroni adjustment (the lowest most significant *P*-value = 0.007 being for the comparison between gene *ORF1ab* and *N*).

There is a much higher number of non-synonymous changes (*n* = 1,293, 62.17%; including missense, start lost and stop gain) compared to synonymous changes (*n* = 787); the ratio non-synonymous/synonymous chances is 0.62. This ratio was also similar when partitioning by main haplogroups; namely, 0.63 for haplogroup A, 0.60 for haplogroup B, and 0.62 for haplogroup A2a (**Table S2**). There is a notable difference between the ratio of non-synonymous/synonymous mutations when analyzed by genes: the maximum value was obtained for gene *ORF6* (0.84), and the minimum for gene *M* (0.41); and the most significant difference was noted between gene *ORF6* and *M*. However, a Fisher’s exact test did not reach significance when adjusted for multiple testing. We also tested for signals of natural selection in SARS-CoV-2 genes by studying *ka*/*ks* index in an interspecific context that includes genomes from pangolin, bat, and SARS coronavirus. For almost all genes, values are below 1, with *ORF1b* being the gene with the lowest value, indicating the action of purifying selection on these genes. Only *ORF10* genes have a positive value, suggesting the action of positive selection operating on this gene (**Table S4**).

We counted 1,731 unique genomes in the database. It is remarkable that a number of sequences were present at high frequencies; for instance, the reference sequence is repeated 78 times; most of them sampled in Asia (76.8%; mostly in China, 60.8%; see below).

Haplotype and nucleotide diversities, as well as Tajima’s *D* test values were computed by geographic regions, haplogroups and genes (**Table S5**). Diversity was very similar in the different continental regions, with very minor differences with respect to the global sample; the only exception is Oceania (represented by Australia and Oceania), which shows notable high diversity values (π = 3.64E-04; *HD* = 9.91E-01). Tajima’s *D* values are significant (below -2) in all regions with the exception of South America.

Diversity indices and Tajima’s *D* statistics are very similar when computed by main haplogroups (**Table S5**). However, diversity values are particularly different when comparing genes. Thus, for instance, *ORF8* shows 23.1 times higher nucleotide diversity (π = 1.07E-03) compared to *E* (π = 8.95E-05); while *ORF1a* shows 48.9 times higher sequence diversity (*HD* = 9.73E-01) than *E* (*HD* = 1.90E-03). *ORF8* and *ORF1a* are the most diverse genes, while *E* shows the lowest diversity. In contrast, Tajima’s *D* values show minor differences between genes.

We investigated mutation variation at two notable sequence features of all SARS-CoV-2 genomes. First, the receptor binding domain (RBD) located in the spike protein, which has been reported to be the most variable part of the coronavirus genome (*5, 6*); these studies indicated that there are six amino acids that are critical for binding to the angiotensin-converting enzyme 2 (*ACE2*) receptor; namely, L455 (pos. 22925-22927), F486 (pos. 23018-23020), Q493 (pos. 23039-23041), S494 (pos. 23042-23044), N501 (pos. 23063-23065), and Y505 (pos. 23075-23077). Second, we also examined variation at the 12 characteristic nucleotide insertion (amino acid sequence PRRA; pos. 23606-23620), which constitutes a polybasic furin cleavage site (PFCS) that is also related to three adjacent predicted O-linked glycans residues: S673 (pos. 23579-23581), T678 (pos. 23594-23596), and S687 (pos. 23621-23623). The low diversity found in these regions is noteworthy. We only found two mutations at the cleavage site (which is present in all the SARS-CoV-2 genomes), namely G23607A (as a non-synonymous change [CGG>CAG or R>Q] private variant in haplogroup A1; GISAID #415709) and 23611G>A (as private synonymous variant of haplogroup A2a2c in GISAID #418390).

We also analyzed the evolution of the diversity indices with time (**Figure S2**; considering only time-points with a minimum accumulated number of 10 sequences). The first genomes sequenced correspond to those sampled in China in late December 2019, followed by genomes from other Asian countries. As expected, sequence and nucleotide diversity experienced an initial rapid growth until 19 January 2020. However, diversity values experienced a very remarkable drop (specially in π; **Figure S2A**) starting on 21 January 2020 and persisting for the next 4 days. Afterward, diversity values progressively grow again from 25 January until reaching the highest values and then a plateau. In non-Asian regions, the pandemic had a delay of a few days (from about 28 January in North America and Europe), but it followed a similarly continuous increase that overtook Asian diversity values. The very high diversity values observed in Oceania are particularly striking (**Figure S2B**) compared to those of other continental regions. Tajima’s *D* values are significantly negative in Asia from the initial outbreak; the growth of this index values is slower in other regions, but it eventually reaches similar values everywhere (see also **Table S5**).

For the beginning of the pandemic, there are only Asian SARS-CoV-2 genomes available in public repositories; however, the number of sequences increases progressively in every region, with Europe being by far the regions that has contributed more genomes to the GISAID database, followed by North America (**Figure S2D**).

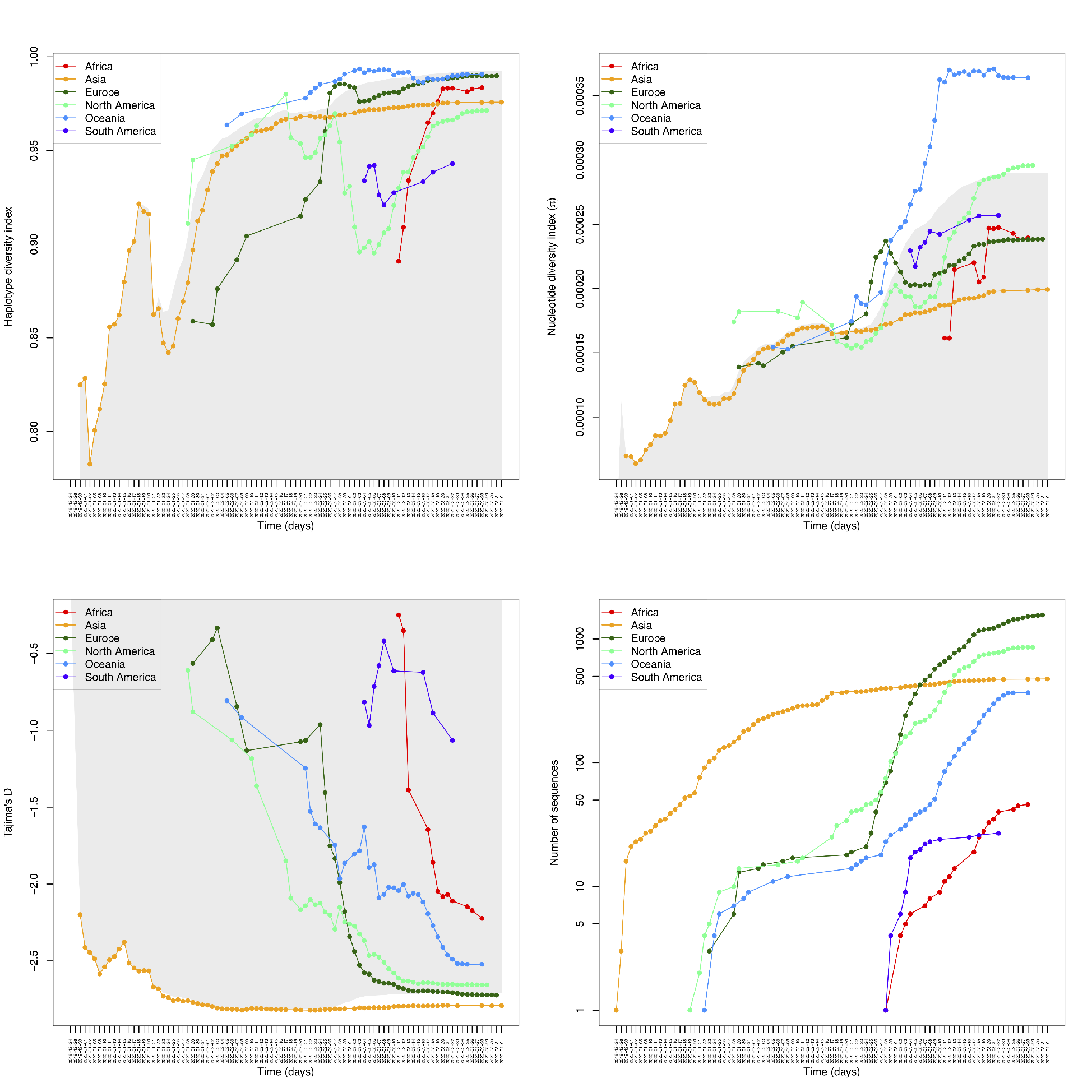

**Figure S2**. Accumulated diversity indices by sampling date in the main regions for the main SARS-CoV-2 phylogenetic branches. The shadowed background area represents the corresponding values for the whole sampling dataset.

**The root of SARS-CoV-2 genomes**

A ML tree was built using pangolin, SARS and bat genomes as outgroups of SARS-CoV-2 genomes (see main text; **Figure S3**). The tree depicts the haplogroup B1 genomes most closely related to bat coronaviruses.

In agreement with the ML tree, we observed that non-human coronavirus sequences from pangolin and bat carry the three transitions C8782T–C18060T–T28144C that identify haplogroup B1 (see below on maximum parsimony tree). There are two exceptions in this alignment. The SARS genome does not present two of the three mutations but this is probably because the SARS sequence has also the lowest identity to SARS-CoV-2, suggesting a relatively long time for the most common ancestor (**Table 1**). As a result, there is a large divergence time (which means plenty of time for reversions) and difficulties for a correct homologous sequence alignment. The genome sequenced from bat #412976 also lacks the three characteristic mutations, but this is most likely due to its very low sequence quality, which can be visually appreciated in the alignment (**Figure S3**) and also in its atypical low identity value to SARS-CoV-2 when compared to the other bat genomes.

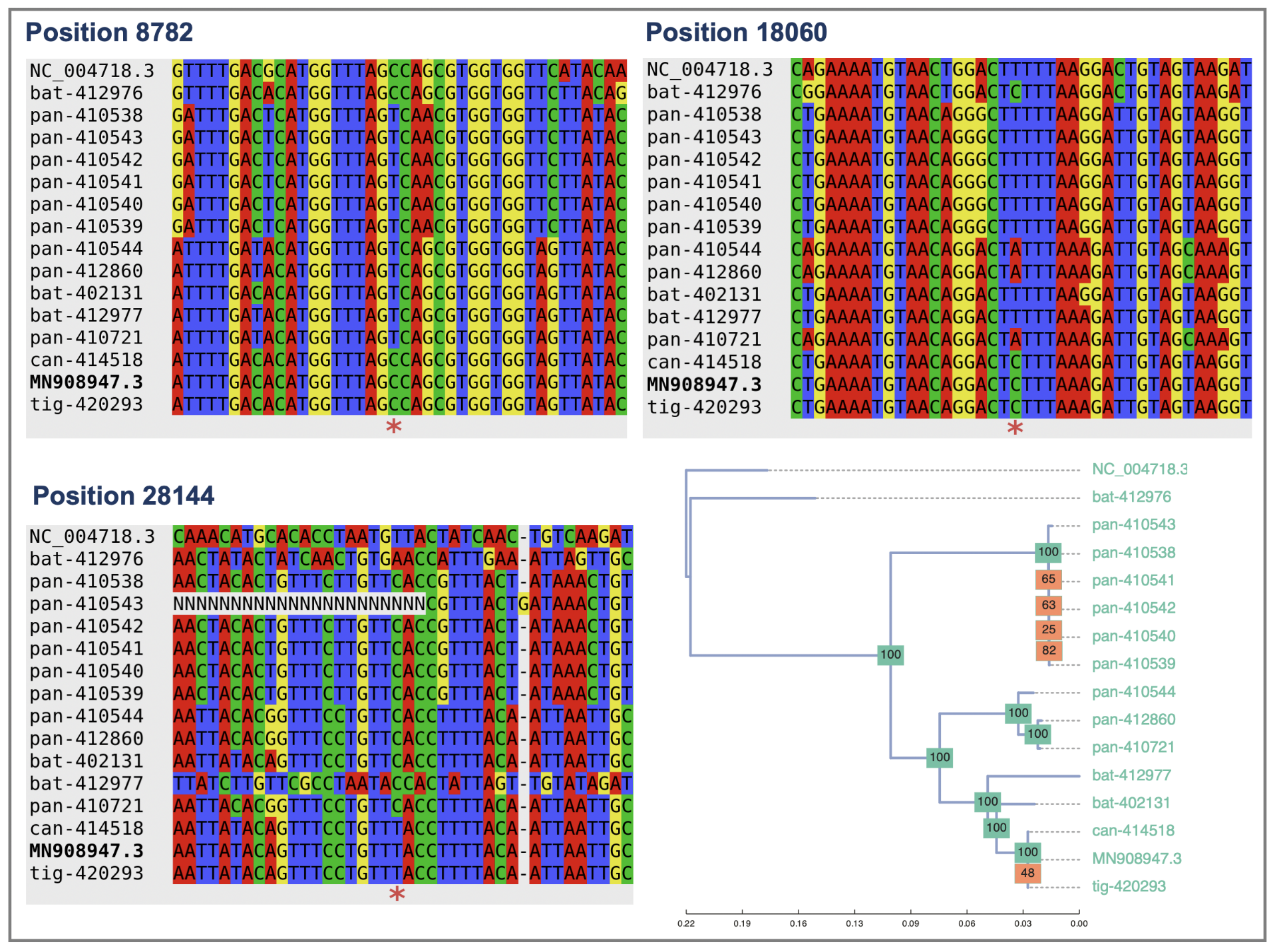

**Figure S3**. Sequence alignments for the segments containing the three variants (stars located at the bottom of the alignments) that characterized the root of all SARS-CoV-2 genomes, and a phylogenetic tree that relates the SARS-CoV-2 human reference (MN908947.3) with other relative coronaviruses in bat, pangolin (“pan”) and SARS (NC_004718.3). We added other human coronavirus segments that were sequenced from an infected tiger and a dog. Alignment is complicated for NC_004718.3 with SARS-CoV-2 due to the reduced identity between them, and this is particularly evident for the segment around position 28144.

**Intraspecific phylogeny of SARS-CoV-2**

We aimed at reconstructing a solid phylogenetic skeleton for SARS-CoV-2 genomes that would allow the identification of diagnostic mutations for main branches. We used a parsimony approach taking advantage of the lessons learnt from the successful human mtDNA tree (*7*); **Figure S4**. A number of studies agree with the definition of two main branches separated by only two transitions: C8782T–T28144C; although different authors use different names for these branches, we follow the labeling initiated by Nextstrain, given its popularity among virologists and other specialists.

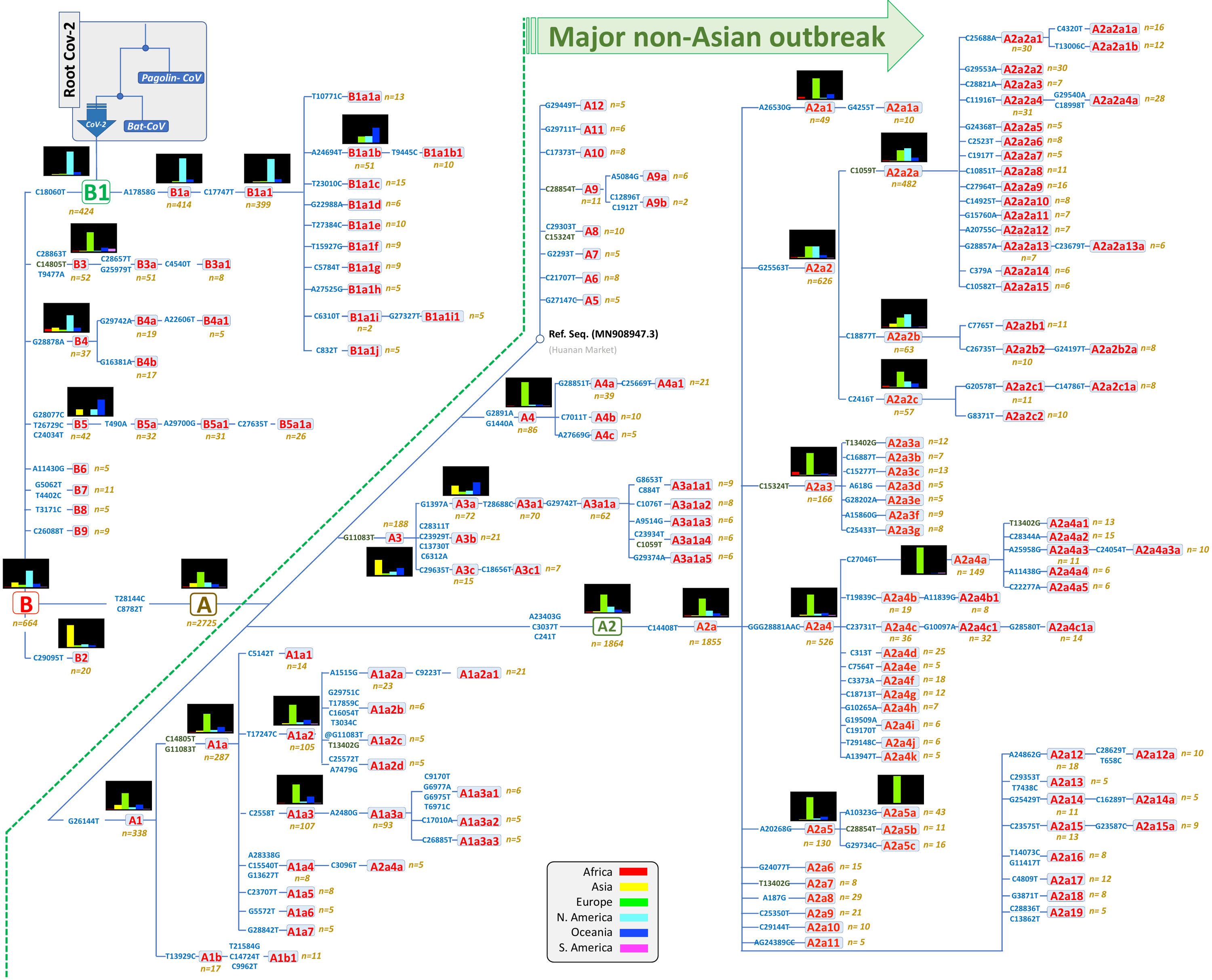

**Figure S4**. Maximum parsimony tree of SARS-CoV-2 genomes. Small histograms represent relative frequencies of the given haplogroup or sub-haplogroup in the different regions. Mutations along branches are referred to changes against the reference sequence. Mutations in dark green color indicate parallel events along the different branches of the phylogeny. Mutations with a @ symbol indicate reversions.

The vast majority of the genomes analyzed in the present study (99.29%) could be unambiguously classified into one of the 164 (sub)haplogroups defined. Only five genomes closely related to the reference sequence could not be classified into either A or B, and there are other 19 genomes that have a problem of resolution but only at the far end of the tree branches. The mutational pathways defining each of these clades are described in **Table S6**.

There number of sequences that have been added to the database increased in a correlated way with the impact of the pandemic worldwide. The main contribution of genomes to GISAID coincides with the non-Asian outbreak, leading to an exponential growth of the number of sequences belonging to haplogroup A (**Figure S5**).

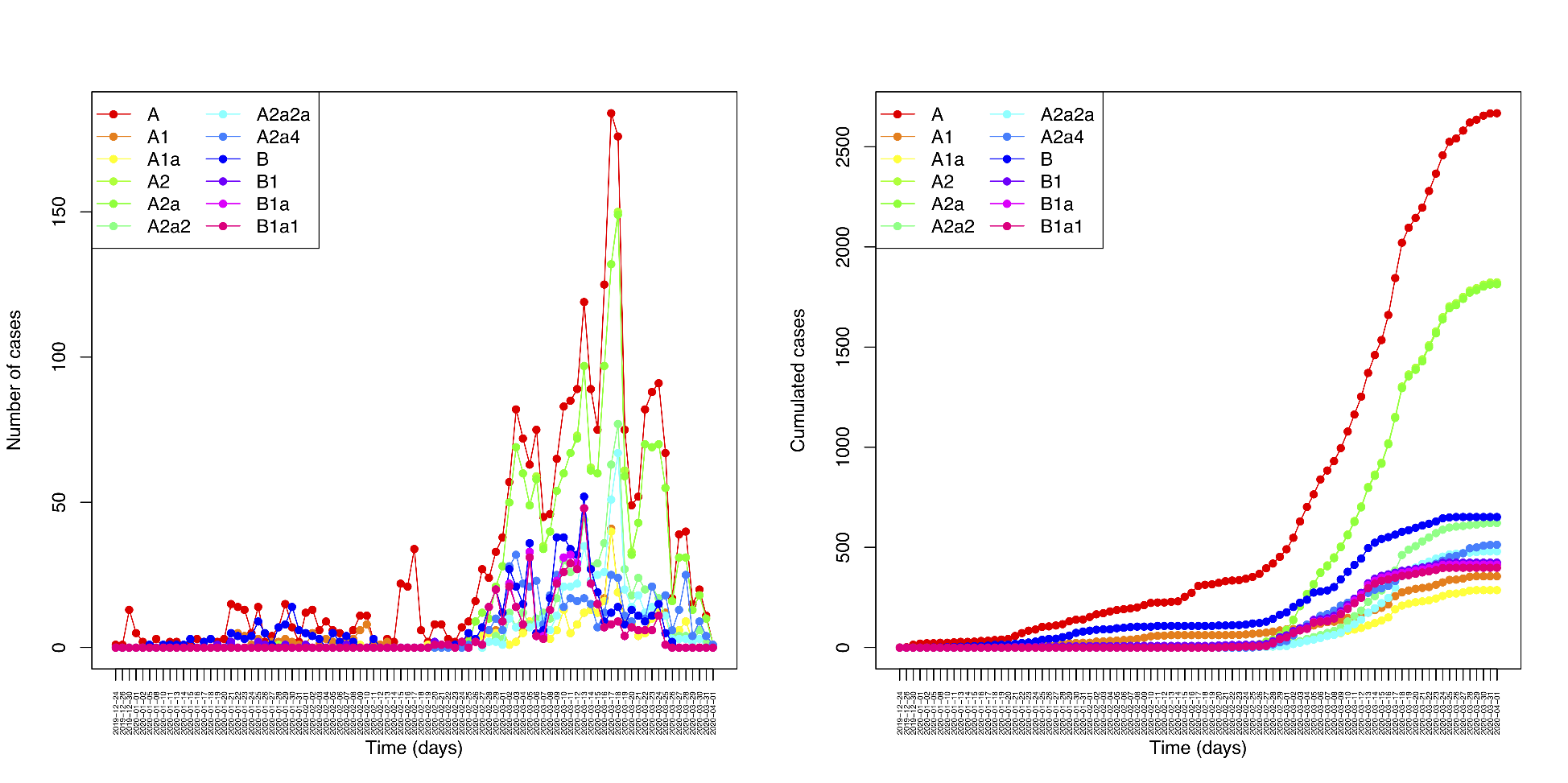

**Figure S5**. Evolution of the number of sequences over time and partitioned by main (sub)haplogroups.

Analysis of diversity indices was also performed on main haplogroups. While π values are almost the same in all regions and haplogroups, HD values are more variable between regions (**Figure S6**).

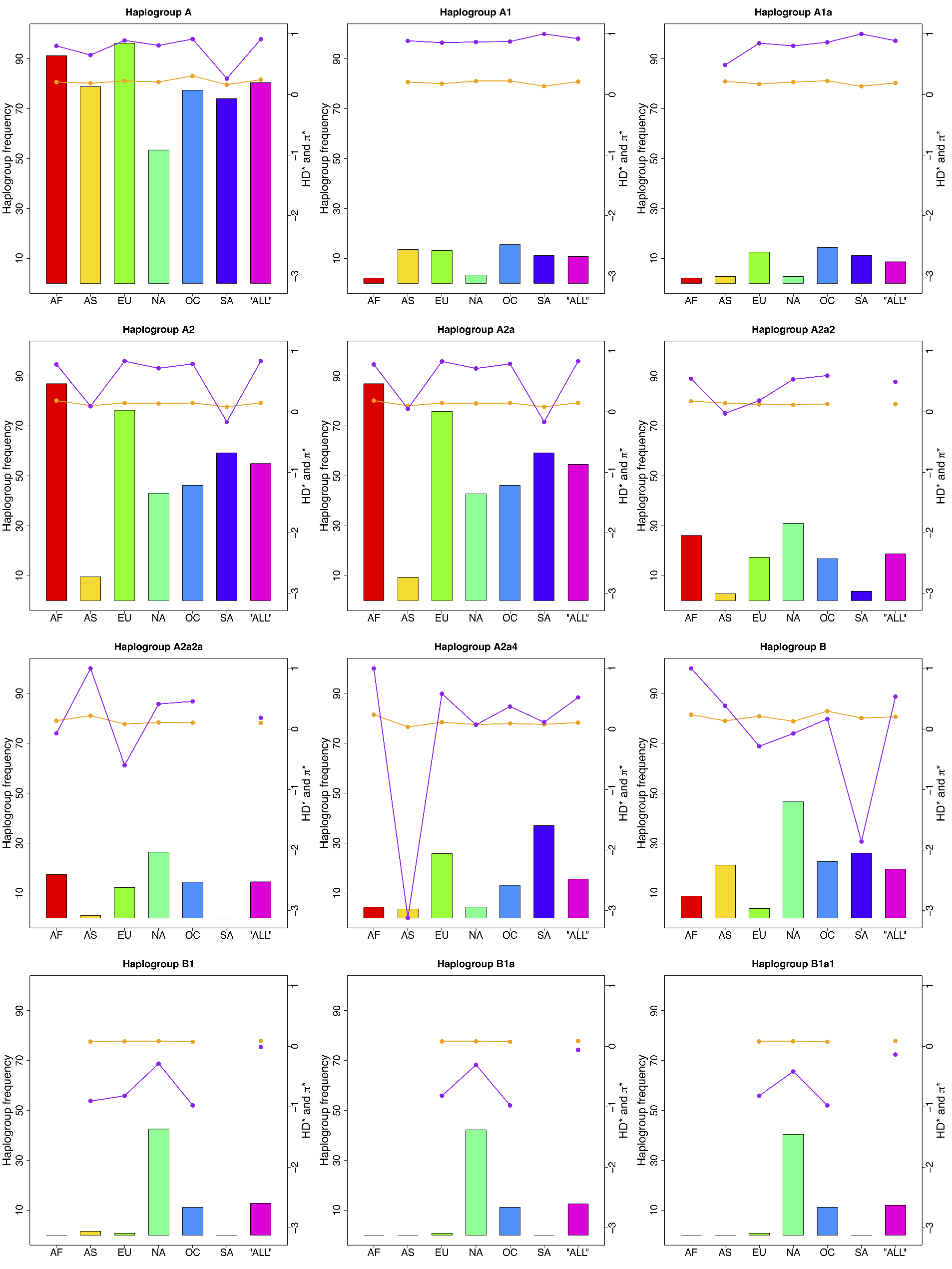

**Figure S6**. Diversity indices computed for main haplogroups. Pink lines connecting dots indicate sequence diversity, and orange lines nucleotide diversity. In order to present both indices together, we depicted values of π/1000 and *HD*×10^–9^.

By counting the occurrence of mutations along the tree branches and those at the tips of the phylogeny (see Methods), it is possible to detect the positions that are mutationally stable from those that are mutational hotspots (**Table S1; Figure S7**). The pattern of occurrences can be summarized as follows:

a) There are 2,147 substitutions (**Table S1**), of which 1,749 (81.46%) occurred only once, and 284 (13.23%) twice in the SARS-CoV-2 genomes; therefore, 94.69% of the mutations occurred no more than twice in SARS-CoV-2 genomes.
b) Mutations along branches in the phylogenetic tree are very stable: there are 185 diagnostic sites in the tree branches (**Table S1**), 110 (59.46%) were singletons, and 154 occurred twice at most (83.24%).
c) There are a few mutational hotspots in the phylogeny; the more important hotspots are C575T (15 hits), 11083 (15 hits), T13402G (13 hits), and A4050C. Note that some rapid mutated mutations, although unstable, can still be diagnostic variants of sub-haplogroups when accompanying other more stable variants (otherwise these changes alone should not be used to define new sub(clades). This is the case of e.g. mutation G11083T with 15 total hits in the phylogeny (three of them accompanying another diagnostic mutation in the tree; this is one of the two most unstable positions in the phylogeny); in combination with the diagnostic positions of A1+C14805T, it is diagnostic for e.g. haplogroup A1a: of the 269 genomes having the sequence motif A1+C14805T, 257 have also G11083T and 19 has only the G11083T mutation.

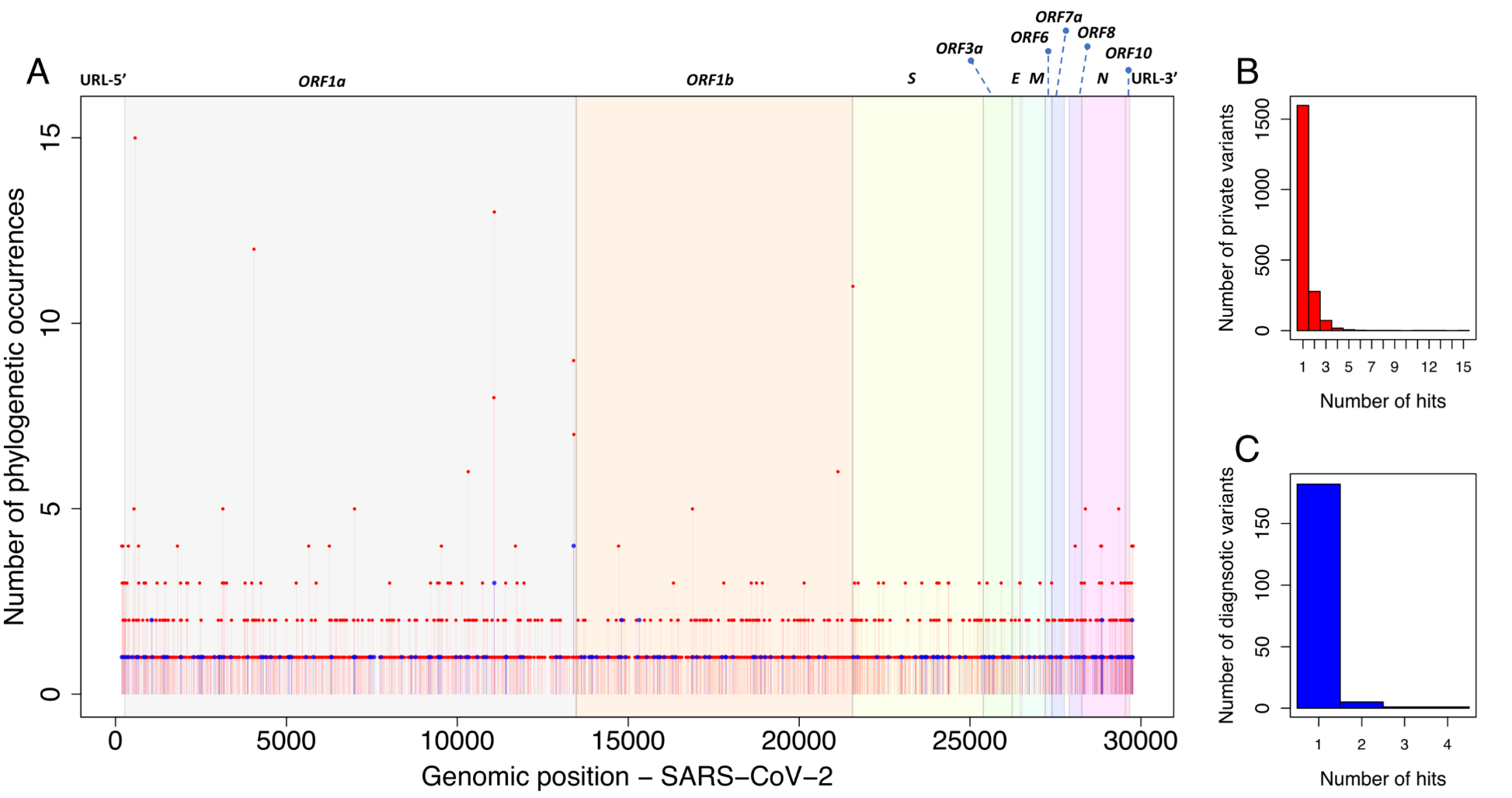

**Figure S7**. (A) Spectra of mutation occurrences along the maximum parsimony tree of **Figure S4**. (B) Number of private variants. (C) Number of mutations located along the branches of the tree (diagnostic variants).

Mutational changes are distributed very homogeneously across the SARS-CoV-2 genomes. In fact, there are very short gaps in the genome of the virus that were not hit by mutations (**Figure S8**).

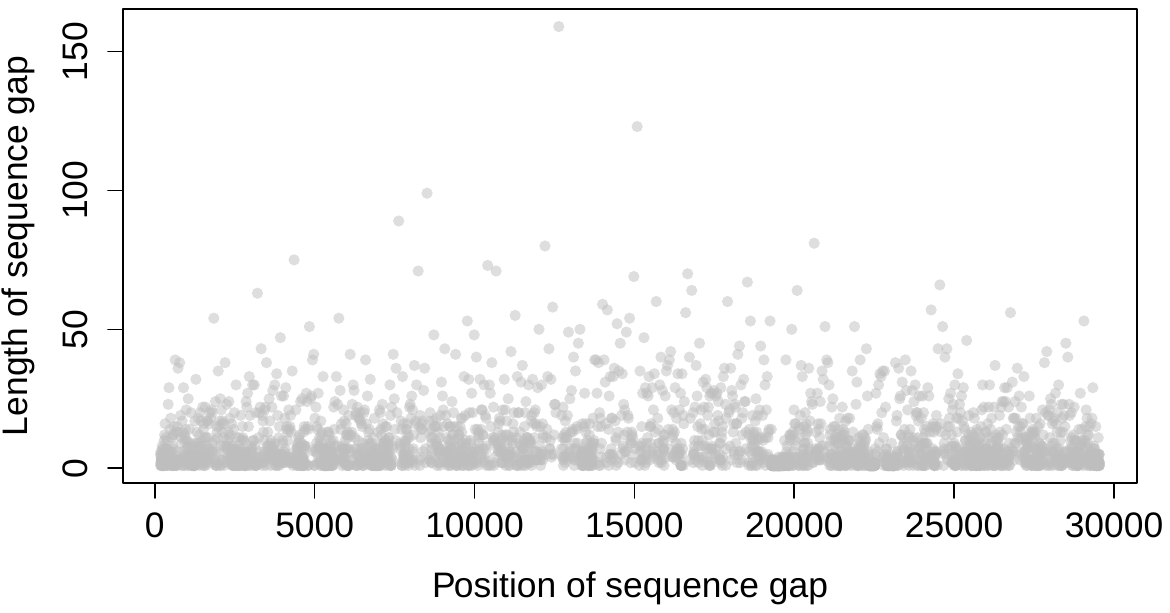

**Figure S8**. Location of the gaps (grey dots) and their length in the genome of SARS-CoV-2 that were not hit by mutations in our dataset.

To further evaluate the robustness of the parsimonious tree, we built a MDS on unique SARS-CoV-2 genomes using the average pairwise discrepancy values for (a) all mutational variants (**Figure S9A**) and (b) only the diagnostic positions of the parsimonious tree (**Figure S9B**). The patterns for both analyses in the MDS plots resemble those inferred from phylogenetic trees.

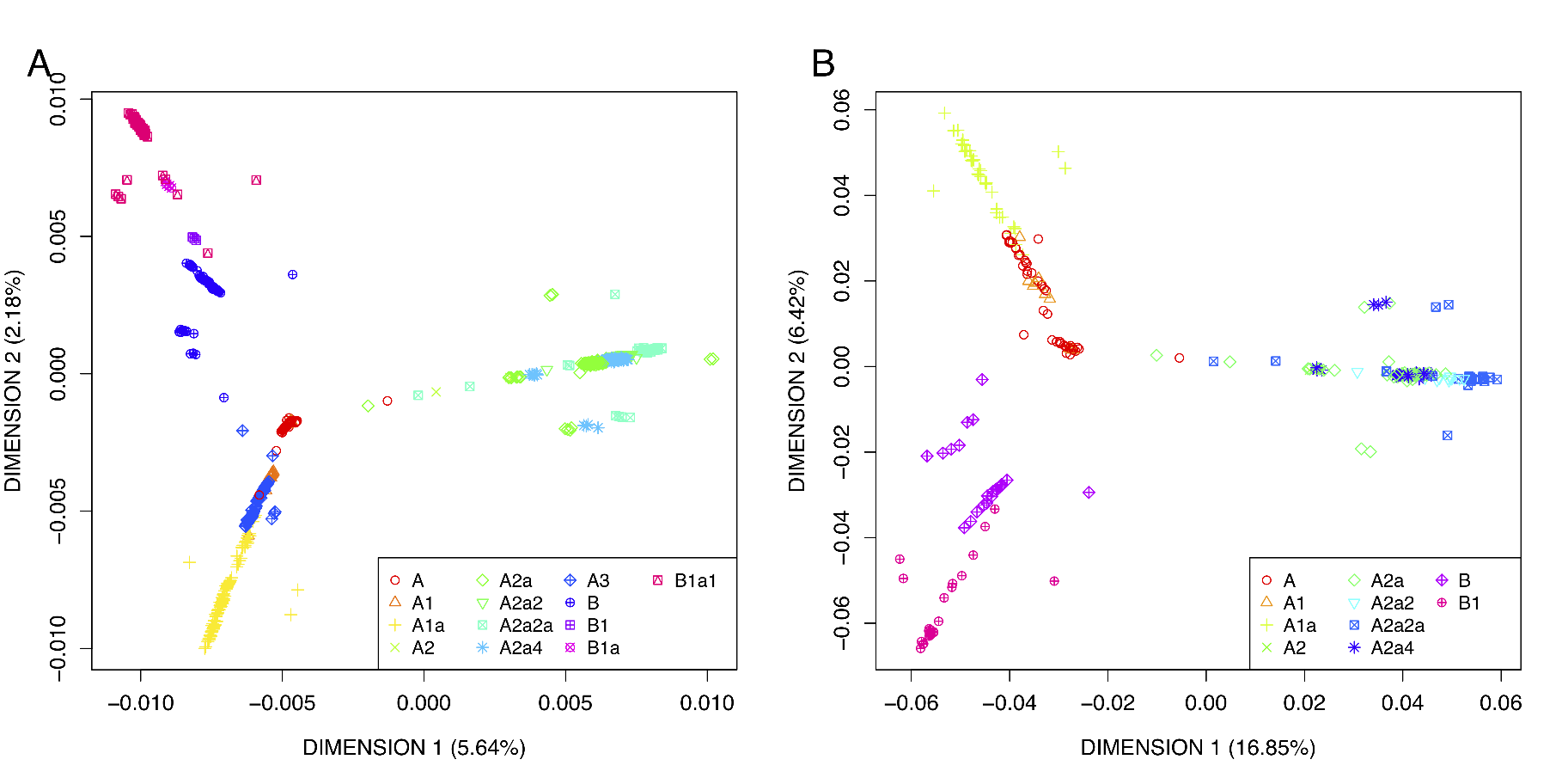

**Figure S9**. MDS plot of SARS-CoV-2 genomes (A) using all mutational variants in the genomes, and (B) using only the diagnostic sites of the phylogeny.

**Considerations on the parsimonious phylogeny**

We built a reference parsimonious phylogeny for SARS-CoV-2 genomes. It reveals that: a) most of the mutations are phylogenetically stable, showing only one or two mutational occurrences in the phylogeny; b) there are only a few mutational hotspots that should not be used to define phylogenetic branches unless accompanied by other more stable mutations; these substitution hotspots do not necessarily relate to the transmission rate of the SARS-CoV-2 and infectivity; c) some mutational instability is detected in some positions located at the tips of the phylogeny; further research is needed in order to determine if this phylogenetic noise is due to mutational instability, sequencing errors or recombination. In this regard, it is important to note that there is an important number of ambiguities in the HQ dataset and that many different sequencing platforms have been used to sequence SARS-CoV-2 genomes, each being prone to specific sequencing artifacts. It can be anticipated that the uniform distribution of mutations along the genome of SARS-CoV-2 will be a challenge for the development and efficiency of future vaccines. Moreover, ML and parsimonious phylogeny suggested that the basal node of B1 is the root of SARS-CoV-2; meaning that the index patient most likely carried this virus strain. Note also that the reference sequence (haplogroup A), does not fall in the root; the decision to select it as the reference does not have a phylogenetic basis.

A practical application of the SARS-CoV-2 tree built in the present study is to facilitate classification of genomes into clades, which might facilitate the work of epidemiologists and other specialists aimed at establishing potential correlations between different clade members and the different clinical phenotypes observed in covid-19, disease severity and differential spread of the disease worldwide. The phylogeny presented is scalable, and nomenclature works in a hierarchical way similar to that demonstrated to be successful in other research areas such as human population genetics (e.g. mtDNA studies).

**Further considerations on phylogeographic patterns of SARS-CoV-2 genomes**

According to GISAID, the first genome to be sequenced was sampled in a patient from China (#402123) on 24 December 2019. A few weeks later, more than a dozen several other genomes had been obtained from Chinese patients from the Hubei province. The first genome sequenced outside Asia corresponds to a sample extracted from a USA patient from Washington (#404895; 19 February 2020). Soon, many other genomes were sequenced from USA, Oceania, Europe, etc. In the database used in the present study, there are genomes sequenced from >62 countries representing the main continental locations: Africa (1.7%), Asia (15.3%), Europe (46.4%), North America (25.3%), and South America (0.6%).

Worldwide patterns of some haplogroup frequencies are summarized in the maps in **Figure S10**.

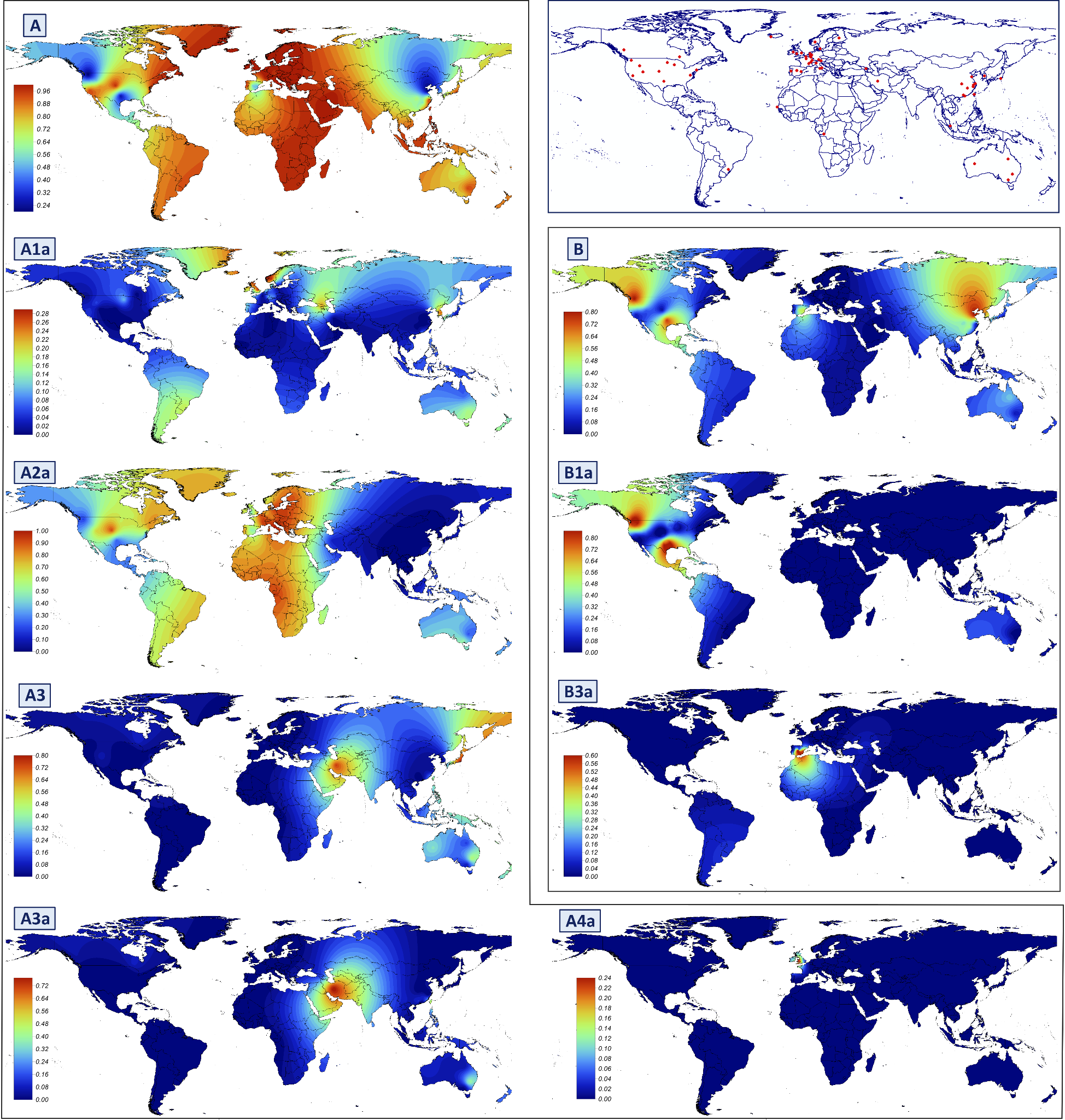

**Figure S10**. Worldwide maps of interpolated haplogroup frequencies. Only the most common (sub)clades are depicted.

**Clade A2 and the main outbreak outside Asia**

Most of the SARS-CoV-2 genomes sequenced between December 2019 the end of February 2020 were sampled from Asian patients. The main SARS-CoV-2 Asian outbreak occurred in January to February 2020. Beginning of March 2020 marked the initial outbreak outside Asia, with clade A2 (and more specifically A2a) being the main responsible for a great proportion of the cases detected worldwide.

Taking advantage of the phylogeny built in the present study, it is possible to infer if the mutations leading to the rise of A2 in Asia or outside Asia. The first A2 genome available in Europe corresponds to a German patient (GISAID: #422425) and was analyzed on 28 February 2020; this genome contains no mutational differences with respect to the root of A2 (**Figure S4**). The next A2 genome sequenced was sampled in a Chinese patient from Shanghai and uploaded to GISAID (#416386) on 31 February 2020. This genome however contains six mutational differences spreading from the root of A2, suggesting that the root of A2 might have been evolving in China weeks or months before February (before the initial Chinese outbreak). Another genome from Asia (Shanghai) was also sequenced a few days later (6 of March 2020) and belongs to the same A2 branch as the previous Chinese one, but with only two mutational differences with respect to the root (**Figure S4**). The first SARS-CoV-2 genome available representing the main phylogenetic sub-branch of A2, namely, A2a (C14408T), corresponds also to a patient from Zheijiang (China) sequenced on 24 February 2020 (GISAID: #422425). The first A2a SARS-CoV-2 European genome corresponds to an Italian sample analyzed on 20 February 2020 (GISAID: #412973). A2a represents the main founder of the non-Asian outbreak; this sub-clade alone accounts for 55.05% of all the genomes available in our dataset (1,853 out of 3,393).

**Considerations on natural selection acting on SARS-CoV-2 genomes**

We explored if patterns of SARS-CoV-2 spread worldwide could be explained by natural selection forces. First, we compared patterns of variation over time in different continental regions (**Figure S2**). Diversity values increased rapidly in China during the first country outbreak. Next, diversity values drop significantly (especially true for the HD) probably coinciding with the role of the ‘super-spreader’ haplotype H4 (reference sequence; see main text) coupled with human intervention in China, which slowed-down further diversification of lineages.

The outbreak outside China to other Asian countries and continents made the curve of cumulative diversity increase to high values of diversity. Again, human intervention worldwide made this exponential increase to reach a plateau. Values of diversity outside Asia overtook those in Asia, because contention of the pandemic by other countries was less efficient than in Asia, as observed from the epidemiological data worldwide. First B, and next A where the main responsible for the initial outbreaks, while A2 provoked the main outbreak outside Asia. The behavior of the curve of Tajima’s *D* index is most paradoxical. Tajima shows very low values from the very beginning and displays the same behavior, but with delay, in the other regions. Significant negative values of this index (below -2) are suggestive of purifying natural selection in DNA sequences; however, negative values could also be compatible with heterogeneity of mutation rates and population expansion, both variables being present in the SARS-CoV-2 pandemic. At the same, human intervention on the disease could also mimic purifying selection.

We further explored the possible action of natural selection by investigating patterns of *ka*/*ks* by genes in groups of SARS-CoV-2 genomes that represent the two extremes of the pathogen genome variation, namely haplogroups B1 and A2a. These values are suggestive of natural selection acting on gene *ORF1a* (ω = 0.19742) and especially on gene *N* (ω = 0.0001); **Table S4**. Only gene *S* shows a moderately positive value (ω = 1.04877), which could be suggestive of slightly positive selection acting on SARS-CoV-2.

The mutational changes differentiating haplogroups A and B do not seem to have relevant pathogenic effects; for instance, C18060T (B1>B), T28144C and C8782T (B>A) are all synonymous changes with low predicted severity (**Table S7**).

Overall, there is suggestive evidence for a role of purifying selection operating during the spread of SARS-CoV-2 worldwide.

**Further considerations on super-spreaders and founder effect**

A total of 48 haplotypes appear at least 7 times in the database (**Table S8**). Almost all of them, with the exception of the reference sequence (H4) that originated at the beginning of the pandemic in China, appear several weeks after the beginning of the Asian pandemic (**Figure S11**), coinciding with the non-Asian outbreak and indicating their important role in the rapid spread of the pandemic.

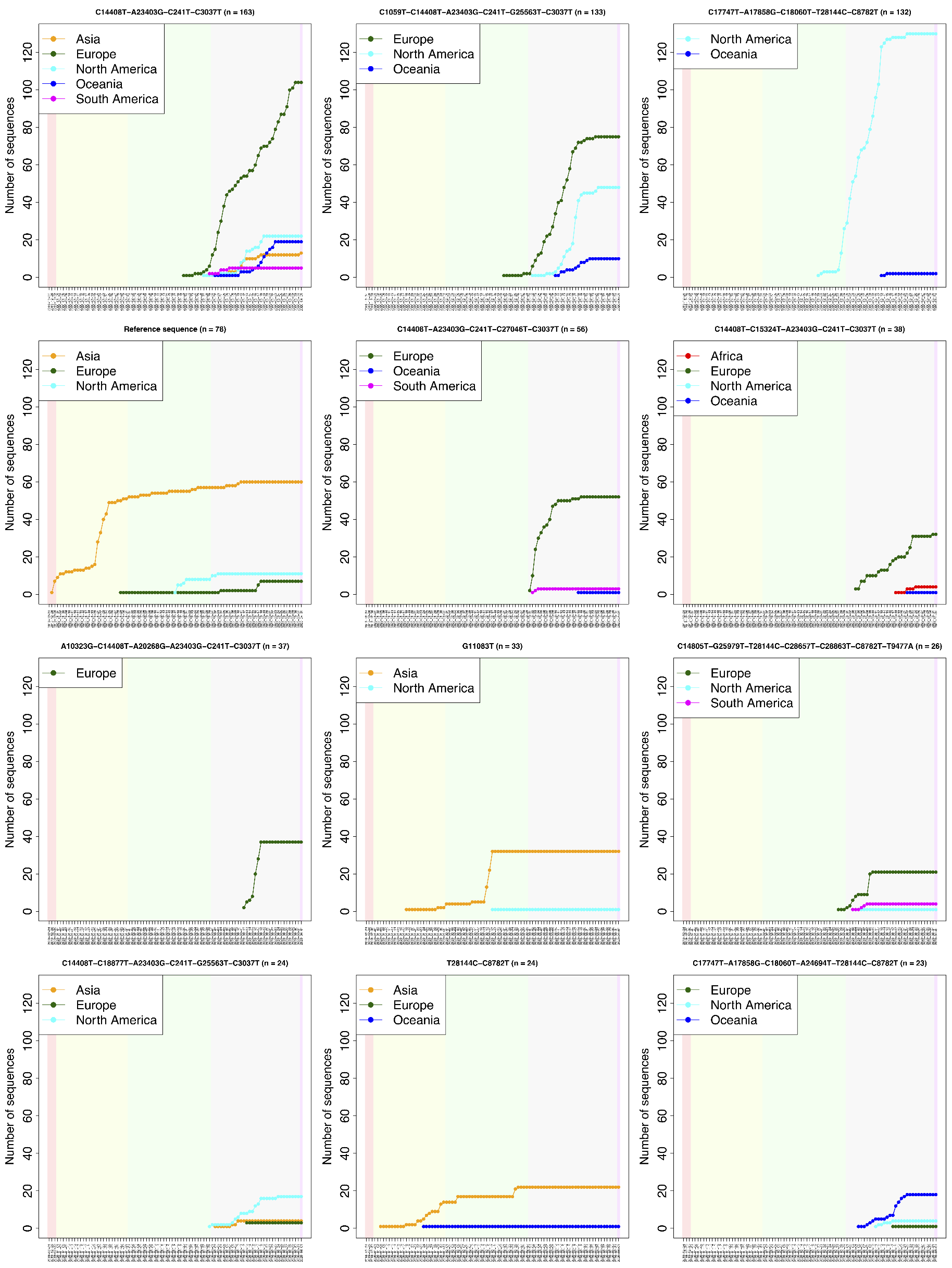

**Figure S11**. Occurrence of the nine most common haplotypes in the SARS-CoV-2 dataset by regions and sampling date.

Network analyses of genome SARS-CoV-2 variation has been carried out for the super-spreading event occurring in the Diamond Princess shipboard (see text for more information) (**Figure S12**).

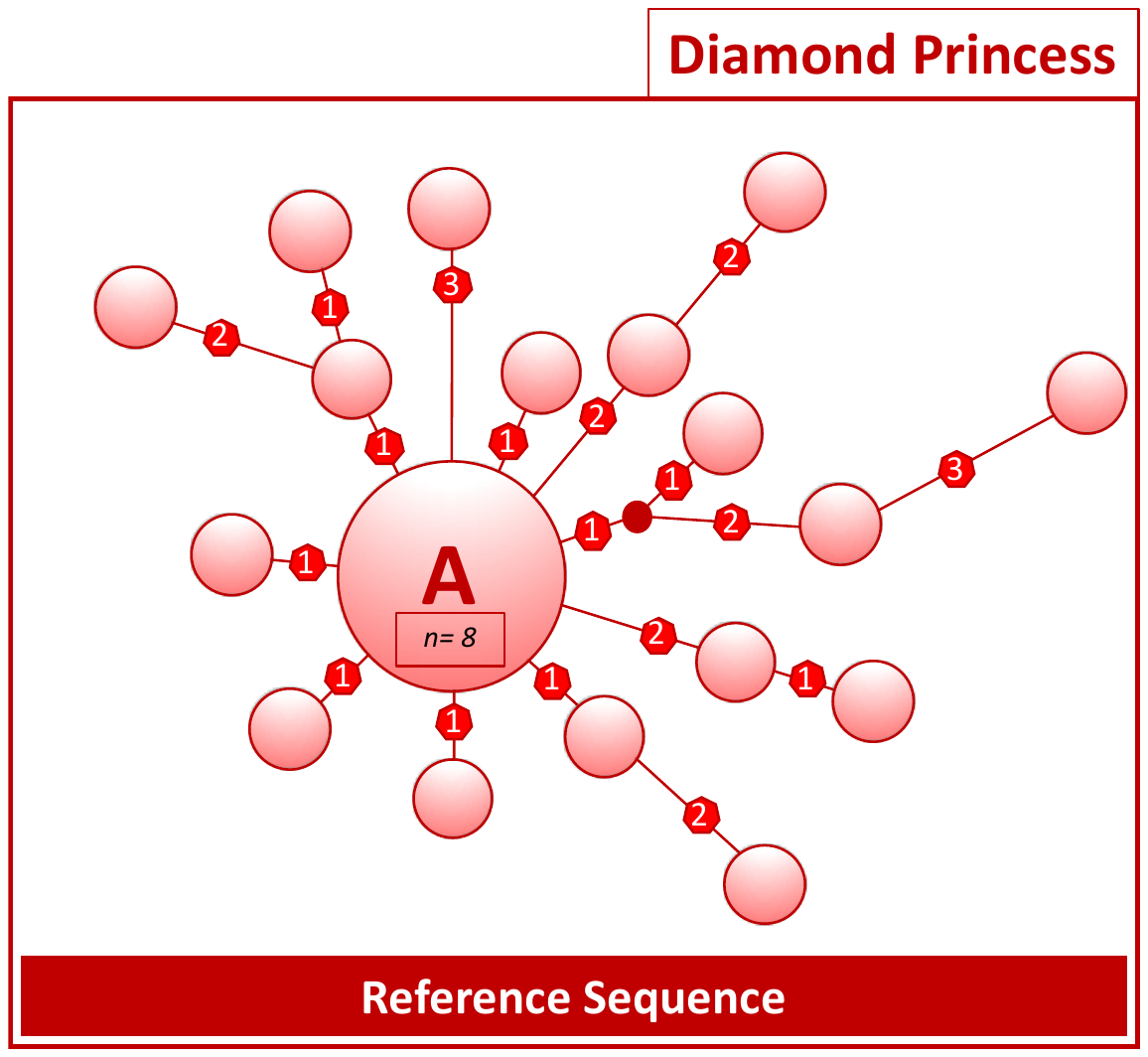

**Figure S12.** Network analysis of the Diamond Princess SARS-CoV-2 genomes. See legend of Figure S1 for further details.

**Association test of haplogroups with sex and age**

Patterns of age and sex were analyzed by regions and main haplogroups (**Figure S13**). The median age was very similar for the different haplogroup categories considered (**Table S9**). The highest female proportion was observed for haplogroup A3 (58.8), and the lowest for B3a (35.4).

Association tests were carried out to evaluate if a particular age group was more severely hit by specific SARS-CoV-2 lineages. An initial test was carried out for the main sub(haplogroups) and age, resulting in two significant associations under a Bonferroni correction, namely, haplogroup A4a (*n* = 39), and B3a (*n* = 51); in both cases, Mann-Whitney test: *P*-value<0.001. Note also that haplogroups A4a and B3a are the ones with the highest median age of all the haplogroups compared but also the largest difference with the median age of the comparable groups (not-A4a and not-B3a). In order to account for the different haplogroup frequency and age patterns existing in each region, we carried out another test considering all the sampling by regions together (*n* = 936); this analysis did not yield significant association (Kruskal-Wallis test: *P*-value = 0.3474).

Additionally, we tested if haplogroups could be related to sex. We found statistically significant association surpassing Bonferroni-corrected significance for haplogroup A3 when compared against non-A3 lineages (*n*_A3_ =y184; Fisher’s Exact Test, *P*-value<0.009).

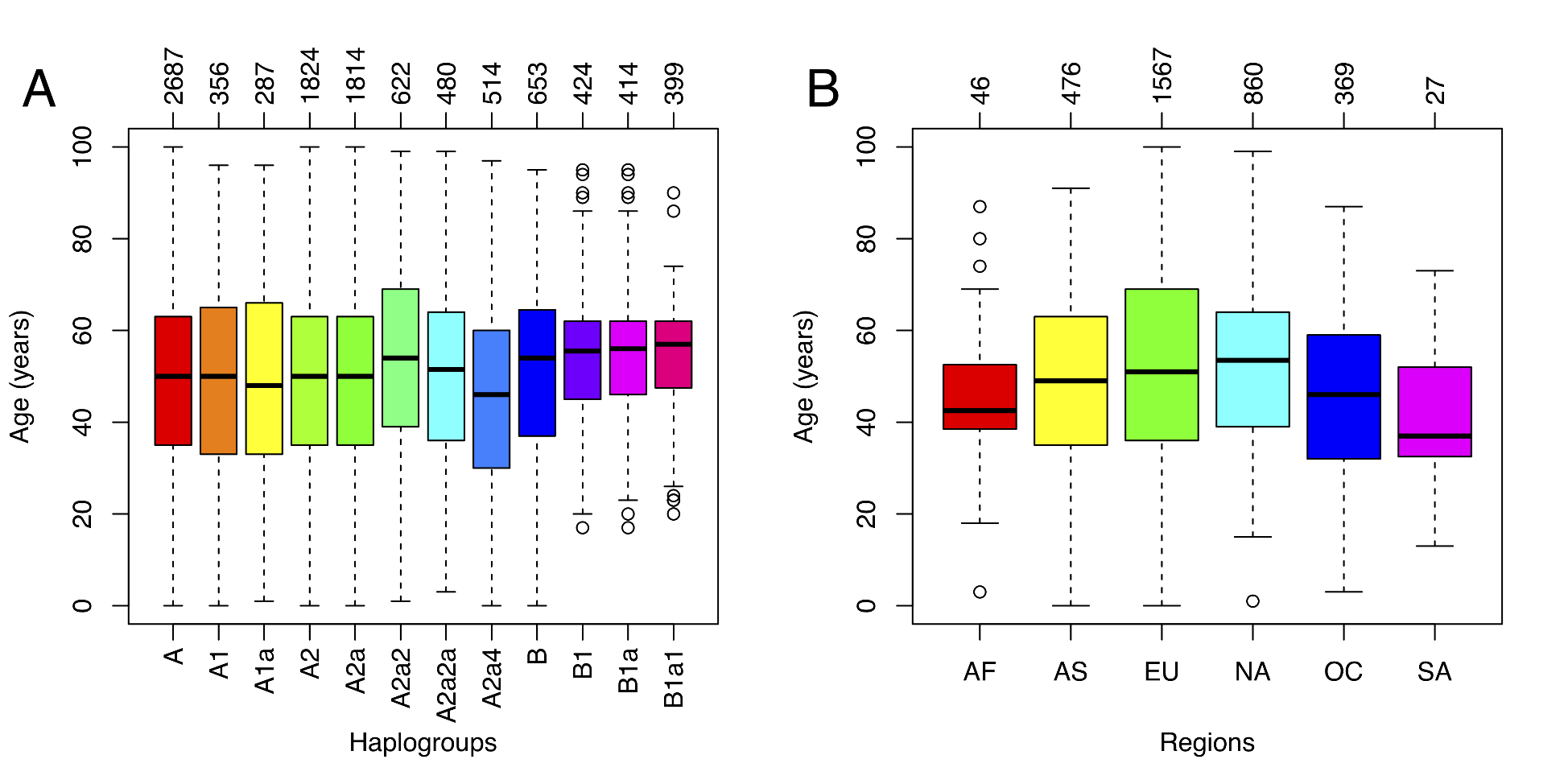

**Figure S13**. Distribution of age by haplogroups and main geographic regions.

**References**

1.	J. A. Rosenfeld, A. K. Malhotra, T. Lencz, Novel multi-nucleotide polymorphisms in the human genome characterized by whole genome and exome sequencing. *Nucleic Acids Res* **38**, 6102-6111 (2010).

2.	I. Ebersberger, D. Metzler, C. Schwarz, S. Paabo, Genomewide comparison of DNA sequences between humans and chimpanzees. *Am J Hum Genet* **70**, 1490-1497 (2002).

3.	M. A. DePristo *et al.*, A framework for variation discovery and genotyping using next-generation DNA sequencing data. *Nat Genet* **43**, 491-498 (2011).

4.	J. M. Zook *et al.*, Integrating human sequence data sets provides a resource of benchmark SNP and indel genotype calls. *Nature biotechnology* **32**, 246-251 (2014).

5.	K. G. Andersen, A. Rambaut, W. I. Lipkin, E. C. Holmes, R. F. Garry, The proximal origin of SARS-CoV-2. *Nat Med* **26**, 450-452 (2020).

6.	Y. Wan, J. Shang, R. Graham, R. S. Baric, F. Li, Receptor Recognition by the Novel Coronavirus from Wuhan: an Analysis Based on Decade-Long Structural Studies of SARS Coronavirus. *Journal of virology* **94**, (2020).

7.	M. van Oven, M. Kayser, Updated comprehensive phylogenetic tree of global human mitochondrial DNA variation. *Hum Mutat* **30**, E386-E394 (2009).
